# FGA139, a Novel Cysteine Protease Inhibitor, Exhibits Anti-Inflammatory and Neuroprotective Activity and Reveals Microglial Modulation via Multi-Omics Profiling

**DOI:** 10.1101/2025.05.03.652036

**Authors:** Ania Canseco-Rodríguez, Florenci V. González, Ana María Sánchez-Pérez

## Abstract

Neuroinflammation is a key driver in the progression of numerous brain disorders, with cysteine proteases such as calpains, caspases, and cathepsins playing central roles in inflammatory signaling.

This study investigates FGA139, a novel irreversible inhibitor targeting cysteine proteases. We evaluated the anti-inflammatory properties of FGA139 in lipopolysaccharide (LPS)-activated macrophages (RAW264.7) and microglia (HMC3). In addition, its neuroprotective effects were assessed in differentiated SH-SY5Y neuron-like cells exposed to conditioned media (CM) derived from the activated immune cells.

FGA139 exhibited a favorable safety profile and robust anti-inflammatory activity, significantly reducing nitric oxide (NO) production in macrophages and TNFα levels in microglia. Conditioned media from both LPS-stimulated immune cells lines (CM^+^) reduced neurite length in neuronal cells. However, CM from HMC3 cells impaired neuronal viability, whereas CM from RAW264.7 cells elevated reactive oxygen species (ROS) and NO levels—indicating distinct neurotoxic signatures. Preincubation of neuron-like cells with FGA139 effectively mitigated most of these adverse effects.

Metabolomic analysis of the activated microglia supernatant revealed that FGA139 increased extracellular levels of neuroprotective metabolites, including purines, linoleic acid, and phenyllactic acid. Proteomic data confirmed that FGA139 attenuated M1-like microglial polarization, likely through modulation of pathways associated with zinc transport and vesicle trafficking.

In conclusion, FGA139 demonstrates potent neuroprotective effects and modulates microglial activation. These findings uncover novel mechanisms underlying the beneficial effects of cysteine protease inhibition and support the therapeutic potential of FGA139 in treating neuroinflammatory conditions, positioning it as a promising modulator of microglial function.

**Highlights:**

- FGA139, an irreversible cysteine protease inhibitor, exhibits anti-inflammatory effects, significantly reducing LPS-induced pro-inflammatory markers (NO in macrophages, TNFα in microglia).
- FGA139 prevented the damaged neurons exposed to conditioned media (CM) from LPS-stimulated immune cells (reduced viability, reduced neurite length and increased ROS/NO).
- FGA139 boosted secretion of neuroprotective metabolites (e.g., purines, linoleic acid) in LPS-stimulated microglia
- Proteome analysis in FGA139 pretreated microglia showed a prevention of LPS-induced M1-like polarization, revealing novel mechanistic pathways.
- FGA139’s ability to modulate microglial activation and protect neurons positions it as a promising candidate for neuroinflammatory diseases.

**Graphical abstract:** 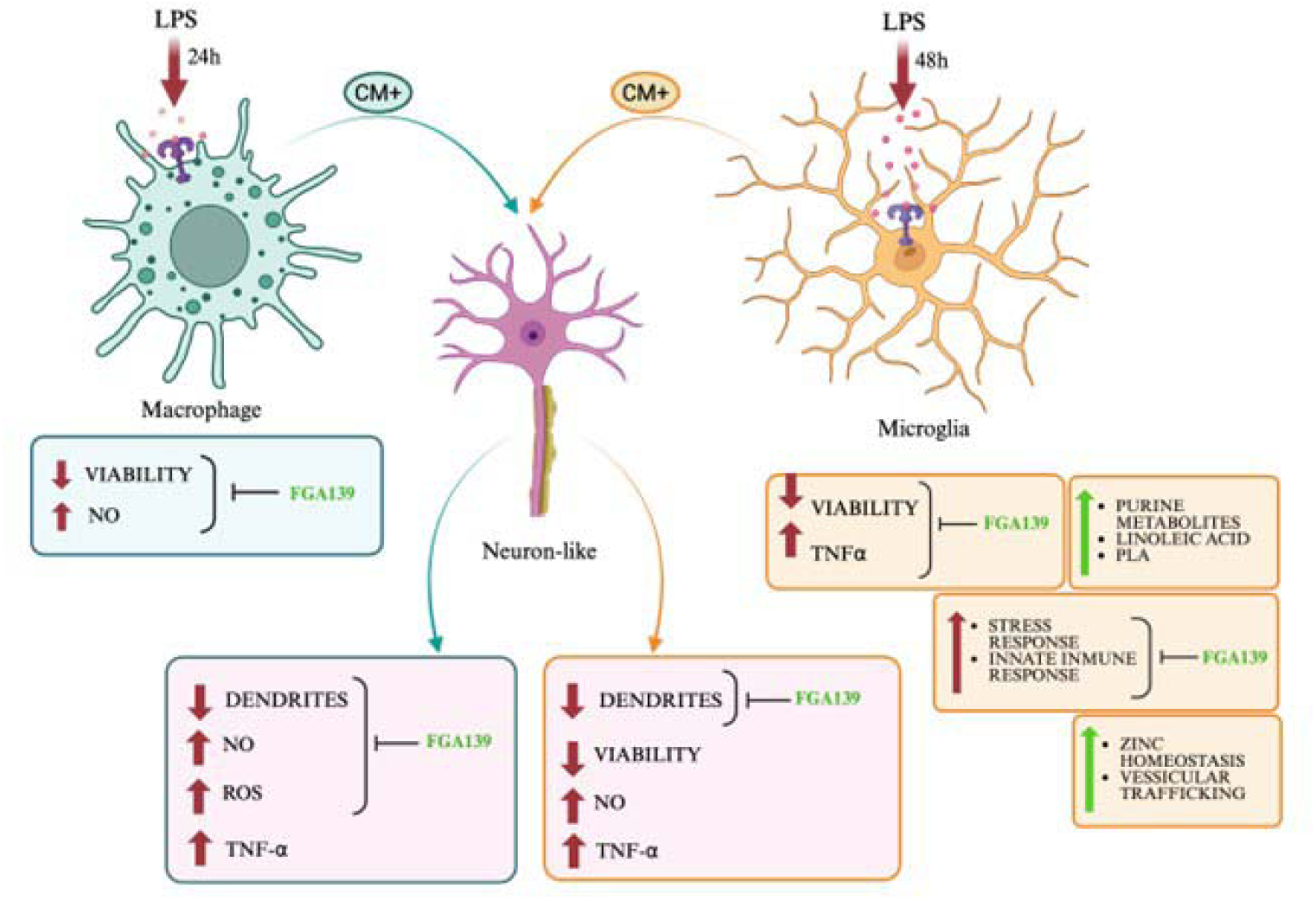

## INTRODUCTION

Neuroinflammation is a critical factor in the pathogenesis of numerous central nervous system (CNS) disorders, including neurodegenerative diseases (Ardura-Fabregat et al., 2017; Sánchez-Sarasúa et al., 2020), psychiatric conditions (Fitton et al., 2022; Meseguer-Beltrán et al., 2023), and traumatic injury (Podbielska et al., 2016). A hallmark of CNS inflammation is the activation of microglia, the resident immune cells of the brain. Sustained microglial activation can promote neuronal damage, which in turn continues microglial reactivity, forming a detrimental feedback loop that leads to neurodegeneration (She et al., 2024; Chiu et al., 2021, Chamera et al., 2020; Podbielska et al., 2016). Also, peripheral macrophages can infiltrate the CNS during inflammatory conditions, like ischemic trauma (Dasari et al., 2021) and age-related neuroinflammation (Wolfe et al., 2018).

Targeting the dysregulated activation of microglia and macrophages has emerged as a promising therapeutic strategy to mitigate neuroinflammation and restore CNS homeostasis (Martínez Leo & Segura Campos, 2020). Among the molecular targets explored, cysteine proteases are of particular interest due to their involvement in key inflammatory and apoptotic pathways (Niemeyer et al., 2020; Pišlar et al., 2021).

Cysteine proteases belong to an extended family of proteases that use cysteine in their active site, but with distinctive function and location. These enzymes include both lysosomal (e.g., cathepsins) and cytoplasmic proteases (e.g., calpains, caspases), which are frequently upregulated in neuroinflammatory contexts (Boland & Campbell, 2004; Islam et al., 2022). For instance, cathepsin B and S (CTSB, and CTSS) contribute to inflammasome activation (Campden & Zhang, 2019,Wendt et al., 2008), while CTSB may also act as a β-secretase, enhancing amyloid burden in Alzheimer’s disease (AD) (Hook et al., 2005). Prolonged activation of calpain further promotes lysosomal rupture and apoptotic signaling (Jeon et al., 2016), and its inhibition has shown neuroprotective effects in models of spinal cord injury (Arataki et al., 2005). These data collectively support cysteine protease inhibition as a rational therapeutic approach in neuroinflammatory conditions (Nakanishi, 2020).

Lipopolysaccharide (LPS), a bacterial endotoxin derived from Gram-negative bacteria, is widely used to model neuroinflammation due to its ability to robustly activate microglia and macrophages via Toll-like receptor 4 (TLR4). This interaction triggers downstream signaling through NF-κB, MAPKs, and inflammasomes, resulting in the release of pro-inflammatory cytokines, reactive oxygen species (ROS), and proteolytic enzymes (Perry et al., 2010; Zhan et al., 2018). Clinically, elevated LPS levels have been detected in both the blood and brain of AD patients (Olsen & Singhrao, 2015). Also gut dysbiosis (Molinero et al., 2023) as well as periodontal bacterial infections (Zhan et al., 2018), characterized by increased Gram-negative bacteria, have been associated with an increased risk of AD pathogenesis. These findings along with animal studies, where LPS administration is used to model neuroinflammation (Skrzypczak-Wiercioch & Sałat, 2022) and impaired cognitive function (Golkar et al., 2025), reinforcing the translational relevance of the LPS model.

In this context, the use of in vitro systems involving LPS-activated microglial or macrophage cell lines provides a valuable platform for investigating molecular mechanisms and screening anti-inflammatory agents. In previous work, we introduced FGA139, a fluorescently labeled, irreversible cysteine protease inhibitor, and demonstrated its protective effects in an in vitro 6-hydroxydopamine (6-OHDA) model of neurotoxicity (Agost-Beltrán et al., 2024).

Building upon these findings, we hypothesize that FGA139, through its irreversible inhibition of cysteine proteases, will attenuate LPS-induced inflammatory responses in immune cells and provide neuroprotection against inflammation-mediated neurotoxicity. To test this hypothesis, we employed two complementary *in vitro* approaches: 1) immune cells (macrophages and microglia) directly stimulated with LPS, where viability, inflammatory markers (NO, TNFα) and reactive oxidative species were measured and 2) differentiated SH-SY5Y neuron-like cells exposed to conditioned media derived from LPS-activated immune cells, where neurite length and cell viability was analysed.

In addition, a comprehensive metabolome and proteome analysis of microglia cells was carried out in microglial cells. By identifying the specific metabolic and proteomic pathways modulated by FGA139 in microglia, this study aimed to uncover novel mechanisms underlying cysteine protease inhibition and its therapeutic potential in neuroinflammatory conditions.

Our findings provide mechanistic insights and highlight FGA139 potential as a therapeutic candidate for neuroinflammation-driven CNS disorders.

## MATHERIALS AND METHODS

### Experimental design

Immune cells, specifically murine macrophages (RAW264.7) and human microglia (HMC3), were seeded. After 24 hours, cells were pre-treated with FGA139 or vehicle control for another 24 hours, followed by incubation with LPS or vehicle for an additional 24 or 48 hours, respectively (**Fig. 1A**). Human neuroblastoma cells (SH-SY5Y) were plated. The differentiation protocol started the following day, for a week. After this period, cells were treated with FGA139 for 24 hours, followed by incubation with conditioned media (CM) from LPS-stimulated RAW264.7 or HMC3 cells for an additional 24 or 48 hours, respectively (**Fig. 1B**).

**Figure 1:**
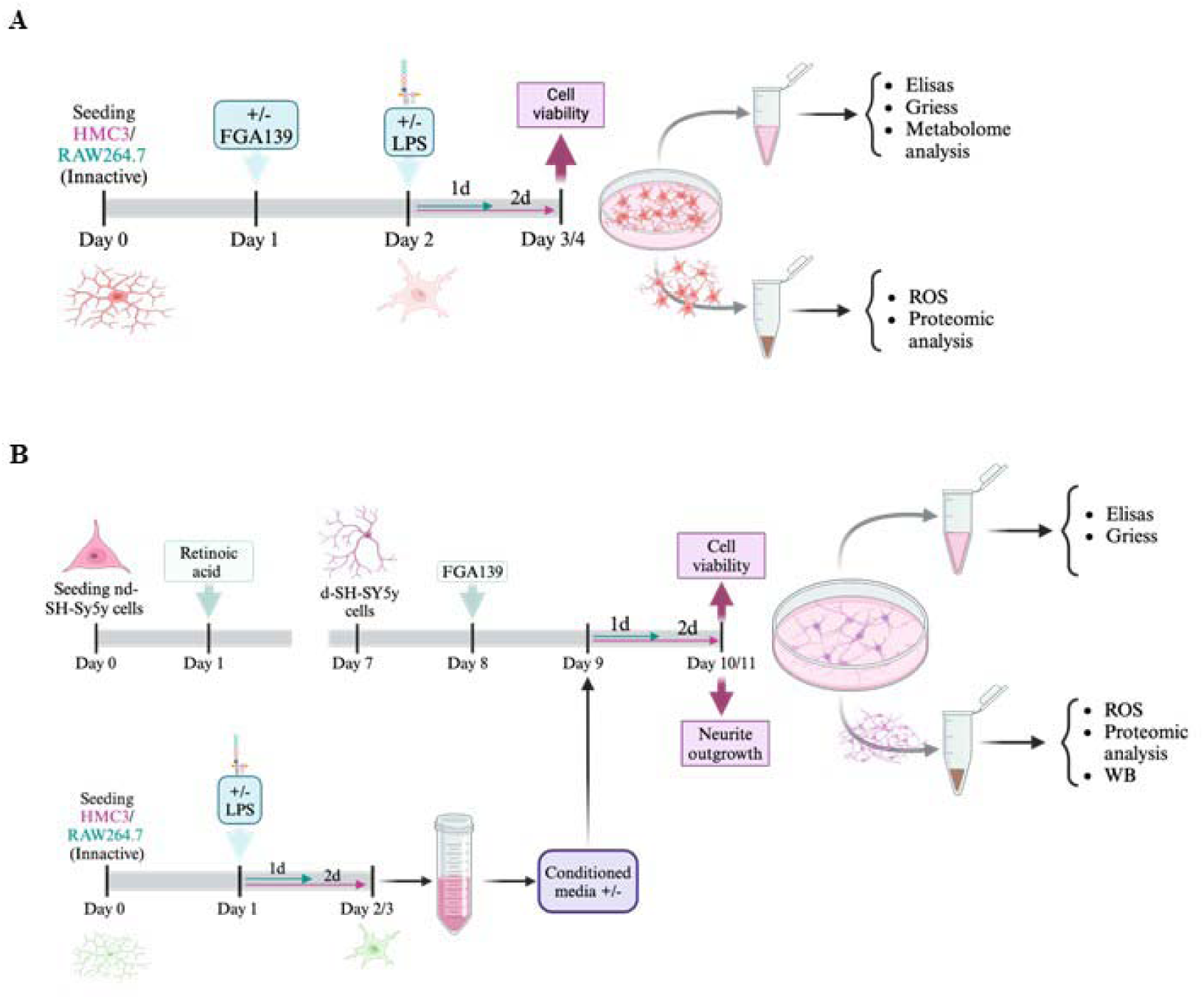
Experimental Design Timeline. **(A)** Human microglia (HMC3) and murine macrophages (RAW264.7) were pre-treated with or without FGA139 before LPS stimulation for 2 and 1 days, respectively. (**B**) Neuroblastoma SH-SY5Y cells were differentiated with retinoic acid (RA) for 7 days, preincubated with or without FGA139, and then exposed to conditioned media from LPS-stimulated immune cells (with their respective LPS-incubation time).

### Cell culture and treatments

Cell lines were cultured in an incubator at 37°C, 5% CO_2_ and 95% humidified air. Cells were maintained in Dulbecco’s Modified Eagle Medium (DMEM; Capricorn, Ebsdorfergrund, Germany), supplemented with 1× L-glutamine (GIBCO, Invitrogen, Paisley, UK), 1× penicillin/streptomycin (GIBCO, Invitrogen, Paisley, UK), and 10% heat-inactivated fetal bovine serum (FBS; Capricorn, Ebsdorfergrund, Germany). When cultures reached approximately 80% confluency, cells were subcultured by washing with phosphate-buffered saline (PBS 1×; Sigma-Aldrich, Velbert, Germany) and detaching with 0.25% Trypsin-EDTA (GIBCO, Invitrogen, Paisley, UK) for 5 minutes at 37°C.

Neuronal differentiation of SH-SY5Y cells was induced by treatment with 10 μM retinoic acid (RA; Sigma-Aldrich, Velbert, Germany) in medium containing 1.5% FBS for 7 days. The medium was refreshed every 3 days.

Stimulation of immune cell lines was achieved by treating cells with lipopolysaccharide (LPS; Thermo Fisher Scientific, Massachusetts, USA) at concentrations of 0.1 or 1 μg/mL. HMC3 cells were stimulated for 48 hours, and RAW264.7 cells for 24 hours. LPS was diluted in Milli-Q water to a final concentration of 1 mg/mL, thoroughly vortexed for homogenization, aliquoted under sterile conditions, and stored at –20°C until use.

All experiments were performed using culture media or cell pellets collected from the distinct cell types (cell passage between 10 and 20).

### Cell viability assay

Cell viability under different conditions was assayed with Alamar Blue following manufacturer’s instructions.

### MAP2 immunostaining and neurite length quantification

SH-SY5Y cells were seeded at a density of 100.000 cells per well on p24 glass coverslips pre-treated with 0.05% PEI (Sigma-Aldrich, Velbert, Germany). After 24 hours, differentiation was induced as described. Following differentiation, cells were treated with 1 or 3 μM FGA139 for 24 hours, followed by exposure to conditioned media (CM) ± LPS (from HMC3 or RAW264.7 cells) for an additional 48- and 24 hours, respectively.

Cells were then fixed with 4% PFA for 15 minutes at room temperature (RT) and permeabilized with 0.3% Triton X-100 in 0.1 M PBS for 15 minutes at RT. Non-specific binding was blocked with 3% BSA for 30 minutes at RT. After several washes, cells were incubated with an anti-MAP2 antibody (1:250, ThermoFisher Scientific, Massachusetts, U.S.). Following additional washes, cells were incubated with a Cy3-conjugated AffiniPure donkey anti-mouse secondary antibody (1:250, Jackson Immuno-Research, Pennsylvania, U.S.) for 1 hour at RT. DAPI (1:2000 in PBS) was then added for nuclear staining.

Coverslips were mounted on glass slides using mounting solution. Fluorescence images were acquired using a Leica DMi8 inverted microscope equipped with a TCS SP8 confocal scanning unit, featuring argon and helium-neon laser beams. Images were captured at excitation and emission wavelengths of 560-586 nm (for the Cy3 secondary antibody) and 358-461 nm (for DAPI). Image analysis was performed using the NeuronJ plugin of ImageJ, measuring the length of primary, secondary, and tertiary dendrites. Images were coded for blinded quantification.

### Western blot analysis

Differentiated SH-SY5Y cells under various conditions were collected, and cell pellets were lysed using RIPA buffer. Protein concentration was measured using the BCA kit (Thermo Fisher Scientific, Massachusetts, USA). Subsequently, 20 μg of protein per sample were separated by sodium dodecyl sulfate–polyacrylamide gel electrophoresis (SDS-PAGE) and transferred to a polyvinylidene difluoride (PVDF) membrane (SERVA Electrophoresis GmbH, Heidelberg, Germany). The membrane was then blocked with 0.3% BSA in TBS-Tween to prevent non-specific binding for 2 hours at room temperature on a shaker. Following blocking, membranes were incubated overnight at 4°C with the following primary antibodies: rabbit anti-pIRS1 S616 (1:500, Invitrogen, Waltham, Massachusetts, USA), rabbit anti-pTAU S396 (1:1000, Invitrogen, Waltham, Massachusetts, USA), and rabbit anti-β-actin (1:1000, Invitrogen, Waltham, Massachusetts, USA). After washing, membranes were incubated with a peroxidase-conjugated anti-rabbit secondary antibody (1:5000, Jackson ImmunoResearch, West Grove, Pennsylvania, USA) for 1 hour at room temperature. Bands were visualized using the ImageQuant LAS 4000 system after thorough washing to remove excess antibodies.

### ELISA, ROS and Griess assays

Extracellular **nitric oxide** (NO_2_ **^-^**) levels were quantified by Griess assay (Promega, Madison, Wisconsin, US) from HMC3, RAW264.7 and d-SH-SY5y cells under the previously described treatments. A reference standard curve was prepared for each assay and added to the plate in duplicates, along with the cell culture media samples, also in duplicates. Once both the kit reagents and the samples reached room temperature, the reagents were added, protected from light, and incubated according to the manufacturer’s instructions. Absorbance was measured at 520 nm within 30 minutes.

Extracellular cytokine levels were measured and quantified by enzyme-linked immunosorbent assay (ELISA) using the culture media from SH-SY5Y, HMC3, and RAW264.7 cells, with or without treatment. After all reagents and samples reached room temperature, standards and samples were added to the ELISA plate in duplicates and incubated according to each assay protocol. Subsequent solutions were added and incubated as specified, with multiple wash steps between each stage. Absorbance was measured after stopping the reactions as follows: for human TNF-α (Thermo Fisher Scientific, Massachusetts, USA), at 450 nm within 2 hours of adding the stop solution; for human IL-1β (Merck, Sigma-Aldrich, Velbert, Germany), immediately at 450 nm; and for human Aβ (Merck, Sigma-Aldrich, Velbert, Germany), at 450 nm and 590 nm within 5 minutes of the final reagent addition, following the manufacturer’s instructions.

Also, HMC3, RAW264.7, and d-SH-SY5Y cells under the indicated treatments were lysed by cold sonication, and centrifuged at 10,000g for 15 minutes at 4°C. The supernatant was collected, and reactive oxygen species (ROS) levels were quantified using the Peroxide Quantification Assay Kit (BQC Redox Technologies, Asturias, Spain) according to the manufacturer’s instructions.

### Metabolomic analysis

Samples were prepared following standard protocols in the analytical unit IIS La FE (Roca et al., 2021). After thawing, 20 μL of each sample was mixed with 180 μL of methanol, vortexed, and incubated for 30min at –20°C. The mixture was centrifuged twice, and 90 μL of the supernatant was combined with 10 μL of internal standard (ISTD; 20 μM mix of caffeine-d9, leucine enkephalin, reserpine, and phenylalanine-d5) for LC-HRMS analysis using a Q Exactive instrument. Quality control (QC) samples were prepared by pooling 10 μL from each sample, and blanks were prepared by replacing biological material with water.

To ensure reproducibility and minimize intra-batch variability, samples were injected in random order. Three QC injections were run at the start for system conditioning (excluded from analysis), followed by QC injections every six samples and a reagent blank at the end.

Chromatographic separation was performed using UPLC coupled to high-resolution Orbitrap mass spectrometry (Q Exactive Plus) under UBMP-standardized conditions for polar compound analysis. Raw data were converted to mzXML format using MSConverter and processed with MEIVEN software and a proprietary UBMP database containing over 400 annotated polar metabolites. Compounds were identified by matching chromatographic peaks to retention times from a validated standard library. Data were analyzed separately for positive and negative ionization modes across three scan ranges, then merged and filtered using QC and blank data. Duplicate features detected in both modes were consolidated, prioritizing those with higher intensity, retention time, and peak quality. The final dataset was normalized using LOESS, log-transformed, and used for downstream statistical analysis.

### Untargeted Metabolomic Data Analysis

Untargeted analysis of polar metabolites was performed by the IIS La Fe Analytical Unit using data previously acquired. Raw data were preprocessed to extract molecular features, defined by a unique combination of mass-to-charge ratio (m/z) and retention time (RT), from both positive and negative ionization modes. After filtering, normalization, and merging of both datasets, a final feature table was generated for statistical analysis.

Differential features between experimental groups were identified using volcano plot analysis followed by orthogonal projections to latent structures (OPLS). Discriminative molecular features were tentatively identified using CeuMassMediator, based on m/z, adducts, and RT, following established metabolite identification levels. Scores generated by CeuMassMediator aided interpretation but were not considered definitive. Inclusion lists were created from selected features for data-dependent acquisition (DDA), and selected QC samples were reanalyzed to acquire MS/MS spectra, aiming to refine identification from level 3 (m/z match) to level 2 (m/z plus MS/MS match). MS/MS spectra were manually compared with CeuMassMediator-integrated spectral databases, as well as MassBank and MassWiki. Features with confident spectral matches were upgraded to level 2 (highlighted in red in the results table, http://hdl.handle.net/10234/730094). Level 2 identifications allowed more precise biological interpretation, though confirmation with analytical standards is still required. For features remaining at level 3, chemical classification (e.g., lipid class) was performed to support broader functional insights and hypothesis generation.

### Proteomic analysis

Proteomic analysis was conducted at the Proteomics Facility of SCSIE, University of Valencia, a member of the ProteoRed network. Protein samples (10µg) were prepared and digested using the SP3 protocol, following standard procedures described by Müller et al. (2019) and Moggridge et al. (2018). Digested peptides were cleaned and processed without further treatment.

For LC-MS/MS analysis, peptides were loaded onto Evotip Pure tips and analyzed using a TimsTOF fleX mass spectrometer (Bruker) coupled with the Evosep One system, using the 30 samples-per-day (SPD) gradient. Peptides were separated on an EvoSep analytical column and analyzed in diaPASEF mode. System performance was monitored using a 50 ng HeLa digest, identifying approximately 6,100 proteins (https://www.evosep.com/wpcontent/uploads/2020/03/Sampleloadingprotocol.pdf). Data were processed using the PASER platform and DIA-NN v1.8 through FragPipe. An in silico spectral library was generated from the SwissProt *Mus musculus* database (17,834 entries). Quantitative analysis employed the DIA-NN Robust LC High Precision workflow, producing summary tables (e.g., combined_protein.tsv) with expression values across all samples.

Further analysis included differential expression analysis, gene ontology enrichment (molecular function, biological process, cellular component), using FragPipe software in TSV format. Protein–protein interactions, functional associations and pathway analysis (e.g., Reactome, KEGG) were explored using the STRING database. Finally, GraphPad software (GraphPad Prism V8 software, GraphPad, La Jolla, CA, USA) was used to visualize the fold changes of significantly differentially expressed proteins between experimental groups, allowing a clearer identification of proteins that were overexpressed or under-expressed under each condition.

### Statistical tests

Data was analyzed using GraphPad software. Data were subjected to the Shapiro–Wilk test for Gaussian distribution. If normality was confirmed, data were represented as mean ± SEM and the “n”, represents the number of independent experiments. One-way ANOVA was carried out, followed by a Tukey’s multiple comparison test to evaluate the effect if CM and the protective effect of FGA139 treatment after CM lesion. Two-way ANOVA was carried out to evaluate the effect of CM lesion and FGA139 treatment on neurite length and was followed by Tukey’s multiple comparisons test. In specific cases, when one variable was compared between two groups, a two-tailed unpaired Student’s T-test with the probability set at < 0.05 was carried out.

## RESULTS

### Protease inhibitor FGA139 rescue cell viability in LPS-stimulated murine macrophages (RAW264.7) and human microglial (HMC3) cells

Initially, three cysteine proteases inhibitors were considered: fluorescent irreversible FGA139, reversible FGA87, and irreversible FGA138. To determine their potential cytotoxicity 1 µM and 5 µM concentrations were tested on murine macrophages RAW264.7. (**Supp. Fig. 1**). FGA139 did not affect viability at any concentration used, hence was chosen for further assays.

As expected, the LPS challenge for 24 hours had a significant impact on RAW264.7 cells (On way ANOVA, F (2,32) = 20.58; *p* < 0.0001). Tukey’s post hoc tests identified significant differences between LPS 0.1 µg/ml (92.07 ± 2.61) and LPS 1 µg/ml (85.92 ± 2.76 %) compared to control (104.1 ± 2.61 %) ([NM vs 0.1 µg/ml LPS], *p* = 0.0127) and ([NM vs 1 µg/ml LPS], *p* < 0.0001), respectively (**Fig. 2A**).

**Figure 2.**
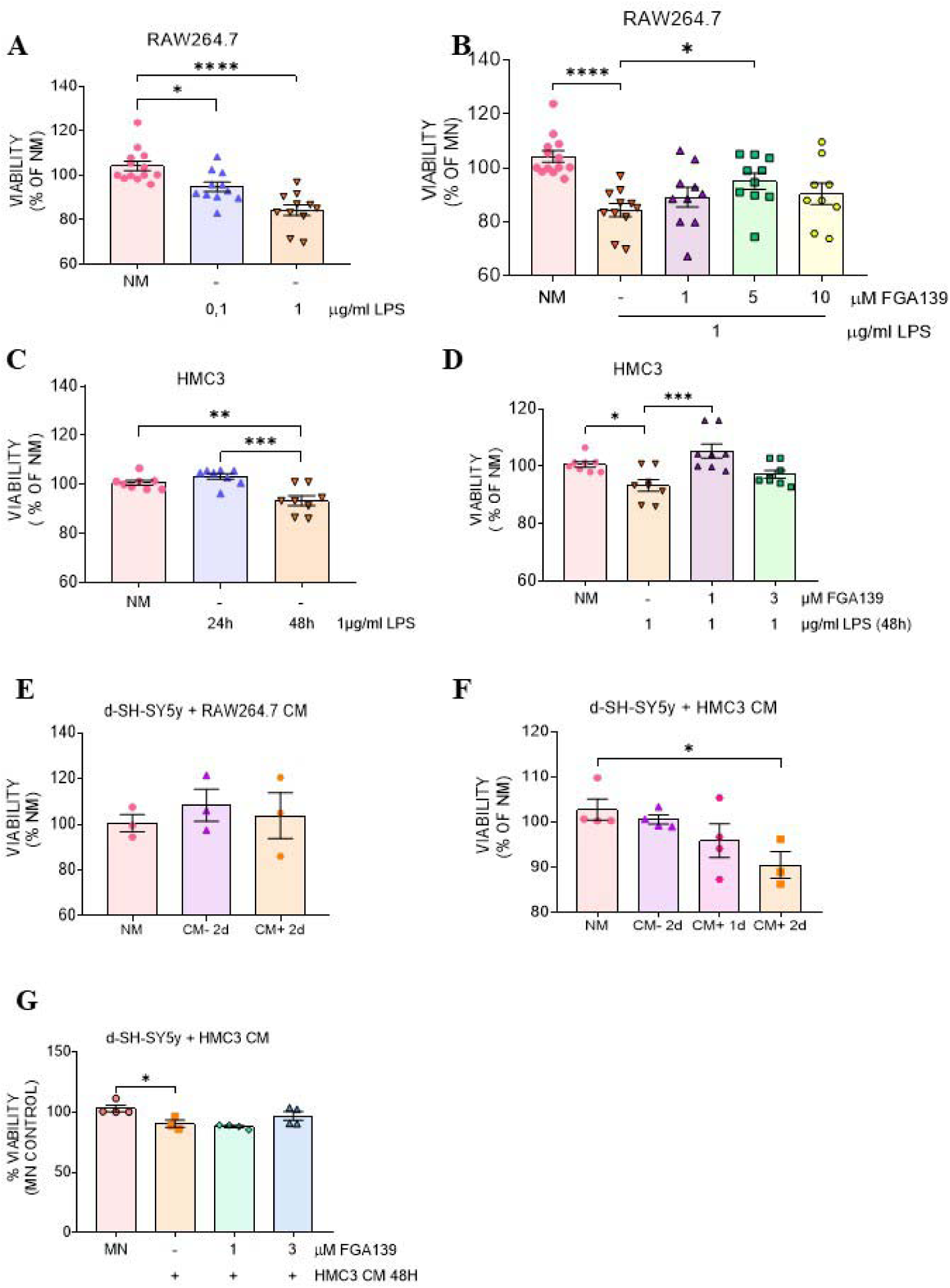
FGA139 Restores Cell Viability in LPS-Treated Immune and Neuronal Cells. (**A**) Viability of RAW264.7 macrophages after 24 h incubation with 0.1 or 1 μg/mL LPS. (**B**) Protective effect of FGA139 on RAW264.7 cells pre-treated for 1 h with 1, 5, or 10 μM FGA139 prior to 24 h exposure to 1 μg/mL LPS. (**C**) Viability of HMC3 microglial cells after 24 or 48 h incubation with 1 μg/mL LPS. (**D**) Protective effect of FGA139 on HMC3 cells pre-treated for 1 h with 1, 5, or 10 μM FGA139 prior to 48 h exposure to 1 μg/mL LPS. (**E**) Effect of conditioned media (CM) from LPS-stimulated and unstimulated RAW264.7 cells on the viability of differentiated SH-SY5Y (d-SH-SY5Y) neuronal cells after 24 or 48 h. (**F**) Effect of CM from LPS-stimulated and unstimulated HMC3 cells on d-SH-SY5Y viability at 24 and 48 h. (**G**) Lack of protective effect of FGA139 on d-SH-SY5Y cells pre-treated with 10 μM FGA139 and subsequently exposed to activated CM from HMC3 for 48 h. Data are presented as mean ± SEM (n = 3-13 per condition). Statistical analysis was performed using one-way ANOVA followed by Tukey’s multiple comparisons test (*p < 0.05; **p < 0.01; ***p < 0.001; ****p < 0.0001).

Based on these results FGA139 (1, 5, 10µM) were used to determine protective activity against LPS (1 µg/ml) on RAW264.7 cells. One-way ANOVA revealed a significant effect of the treatment (F (2,31) = 17; *p* < 0.0001). Tukey’s post hoc tests analysis identified that cell viability improvement by 5 µM FGA139 (95.03 ± 2.99 %) was significantly higher than in LPS alone (85.92 ± 2.76 %) ([LPS vs 5 µM FGA139 + LPS], *p* = 0.0159) (**Fig. 2B**).

Similarly, we assessed the viability of human microglial HMC3 cells following treatment with 1 µg/mL LPS for 24 or 48 hours. One-way ANOVA revealed a significant effect of treatment, F (2, 21) = 12.73, *p* = 0.0002. Tukey’s post hoc test indicated that after 48 hours, cell viability significantly decreased from 100.7 ± 0.74% to 93.33 ± 2.21% ([NM vs. LPS 48 h], *p* = 0.0046) (Fig. 2C). (**Fig. 2C**).

To assess the protective effect of FGA139, we tested its efficacy against the 48-hour LPS treatment. One-way ANOVA showed a significant effect, F (2, 21) = 10.10, *p* = 0.0008. Tukey’s post hoc test revealed that pre-treatment with 1 µM FGA139 significantly restored cell viability, increasing it to 105.4 ± 2.46% ([LPS vs. 1 µM FGA139 + LPS], *p* = 0.0006) (**Fig. 2D**).

### Conditioned media (CM) from activated HMC3, but not from RAW264.7 reduce differentiated SHSY5y cell viability

We next investigated whether conditioned media from LPS-stimulated immune cells (CM^+^) affected the viability of d-SH-SY5Y cells. One-way ANOVA indicated no significant effect of treatment with CM from RAW264.7 cells (24 h and 48 h), as d-SH-SY5Y cell viability remained unchanged (**Fig. 2E**). In contrast, treatment with CM from HMC3 cells for 48 h significantly reduced d-SH-SY5Y cell viability from 102.7 ± 2.34% to 90.92 ± 2.43%. One-way ANOVA showed a significant effect, F (3, 11) = 3.75, *p* = 0.0447, and Tukey’s post hoc test confirmed a significant reduction in viability ([NM vs. HMC3 CM], *p* = 0.0438) (**Fig. 2F**).

To explore the potential neuroprotective effects of cysteine protease inhibitors, we first evaluated the impact of FGA139, FGA138, and FGA87 on d-SH-SY5Y cell viability, as previously done for immune cells (**Supp. Fig. 1B**). FGA139 was found to be safe at concentrations of 1 and 3 µM. However, pre-treatment with FGA139 did not significantly improve d-SH-SY5Y cell viability following exposure to HMC3-derived CM . One-way ANOVA showed no significant treatment effect (**Fig. 2G**), indicating a lack of protective action under these conditions.

### Conditioned media (CM) from LPS-activated RAW264.7 and HMC3 cell lines reduced dendrite length in d-SH-SY5Y cells, and preincubation with protease inhibitors counteracted this effect

To assess the impact of CM^+^ from stimulated RAW264.7 cells and HMC3 on d-SH-SY5Y cells, dendrite length was analyzed using MAP2 immunostaining. Representative images are shown in **Fig. 3A**. Two-way ANOVA revealed a significant main effect of FGA139 treatment (F (2,31) = 9.217, *p* = 0.0007), as well as a significant interaction between CM and FGA139 (F (2,31) = 14.87, *p* < 0.0001), indicating that RAW264.7 CM only reduced dendritic length in the absence of FGA139. Specifically, CM significantly reduced mean primary dendrite length from 84.14 ± 3.47 µm to 53.52 ± 2.35 µm ([NM vs. CM], *p* < 0.0001), while pre-incubation with 1 µM FGA139 prevented this reduction (85.52 ± 7.89 µm, [CM vs. 1 µM FGA139 + CM], *p* = 0.0008) (**Fig. 3B**).

**Figure 3.**
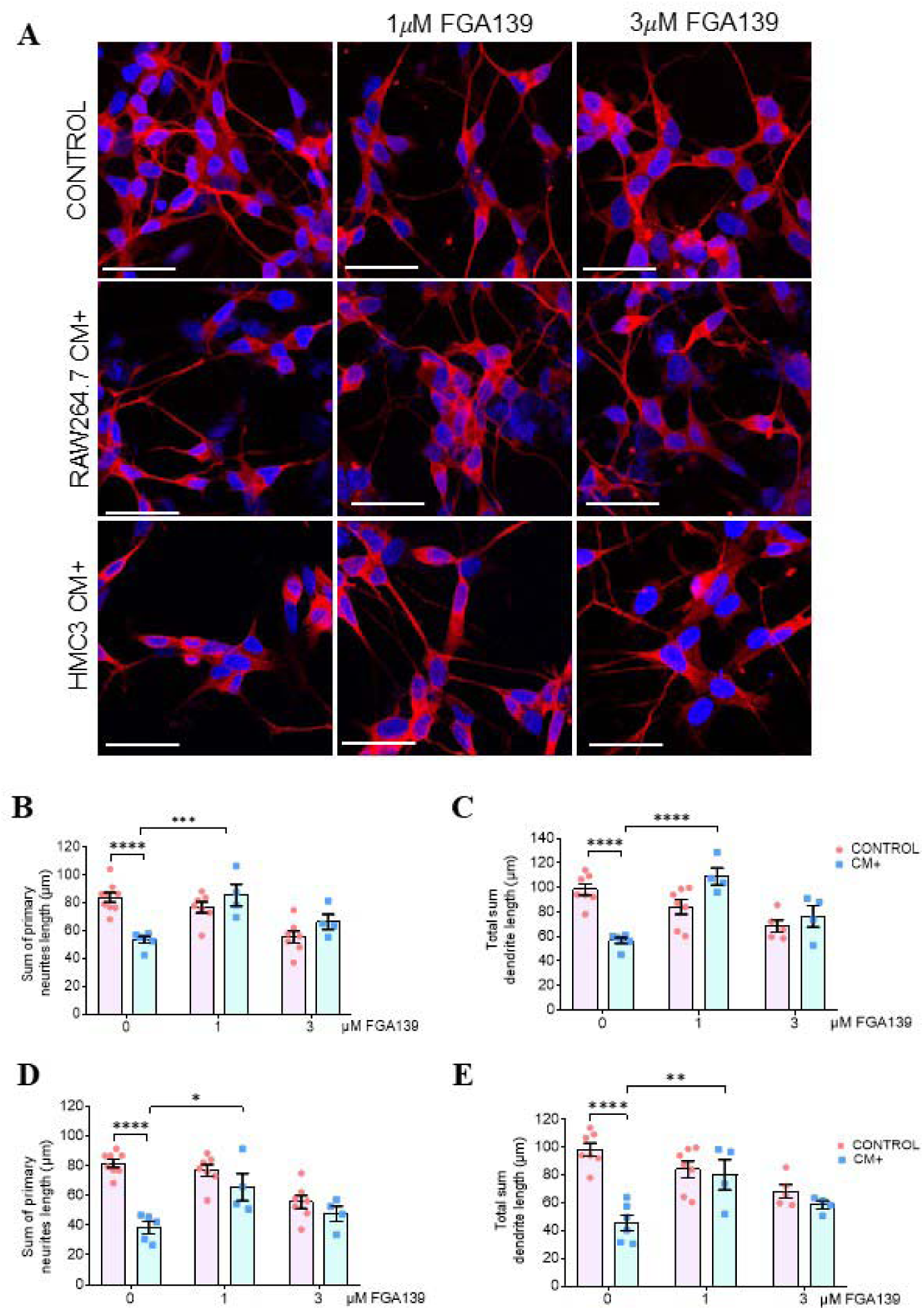
FGA139 Preserves Dendrite Length in d-SH-SY5Y Cells Exposed to Conditioned Media from Stimulated RAW264.7 and HMC3 Cells. (**A**) Representative immunofluorescence images of differentiated SH-SY5Y cells stained with anti-MAP2 (red), pre-treated with FGA139 for 24 h before incubation with conditioned media (CM) from RAW264.7 (24 h) or HMC3 (48 h). (**B**) Quantification of primary neurite length and (**C**) total dendrite length in cells exposed to RAW264.7 CM. (**D**) Quantification of primary neurite length and **(E**) total dendrite length in cells exposed to HMC3 CM. Data are presented as mean ± SEM (n = 4-8 per condition) and analyzed by two-way ANOVA followed by Tukey’s multiple comparisons test (*p < 0.05; **p < 0.01; ***p < 0.001; ****p < 0.0001). Scale bar: 36.8 μm.

Similarly, total dendrite length analysis (sum of primary, secondary and tertiary dendrites) was significantly affected by FGA139 (F (2,27) = 9.493, *p* = 0.0008), with a significant interaction also observed (F (2,27) = 20.43, *p* < 0.0001). CM significantly decreased total dendrite length from 98.46 ± 4.69 µm to 56.67 ± 2.39 µm ([NM vs. CM], *p* < 0.0001), whereas 1 µM FGA139 effectively prevented this reduction (109.00 ± 6.95 µm, [CM vs. 1 µM FGA139 + CM], *p* < 0.0001) (**Fig. 3C**).

To further investigate the neuroprotective potential of FGA139, d-SH-SY5Y cells were exposed to HMC3 CM with or without FGA139 pre-treatment. Two-way ANOVA showed a significant main effect of FGA139 on primary dendrite length (F (2,28) = 5.384, *p* = 0.0105), as well as a significant interaction with CM (F (2,28) = 7.103, *p* = 0.0032).

CM significantly reduced primary dendrite length from 81.60 ± 2.68 µm to 38.35 ± 4.09 µm ([NM vs. CM], *p* < 0.0001), while 1 µM FGA139 significantly mitigated this effect (65.15 ± 9.15 µm, [CM vs. 1 µM FGA139 + CM], *p* = 0.0102) (**Fig. 3D**).

Total dendrite length was also significantly affected (F (2,27) = 4.182, *p* = 0.0262), with a notable interaction (F (2,27) = 10.25, *p* = 0.0005). CM reduced total dendrite length from 98.46 ± 4.69 µm to 45.76 ± 5.63 µm ([NM vs. CM], *p* < 0.0001), while 1 µM FGA139 significantly preserved dendritic structure (80.37 ± 10.98 µm, [CM vs. 1 µM FGA139 + CM], *p* = 0.0093) (**Fig. 3E)**.

Interestingly, treatment with 3 µM FGA139 under normal conditions (without CM) led to a reduction in primary dendrite length (68.44 ± 4.95 µm, [NM vs. 3 µM FGA139], *p* = 0.0001), suggesting that while FGA139 offers protection under inflammatory conditions, its inhibition of protease activity under basal conditions may be detrimental. This finding supports the hypothesis that cysteine protease activity is elevated under inflammatory conditions, and its selective inhibition represents a viable neuroprotective strategy.

### CM from LPS-activated RAW264.7 and HMC3 cell lines does not alter IRS1 ord Tau phosphorylation

To explore the mechanisms underlying the effects of CM on d-SH-SY5Y cells, we assessed the phosphorylation levels of pTau (S396) and pIRS1 (S616), two well-established markers of neurodegeneration (Petrozziello et al., 2022). Western blot analysis revealed no significant changes in the phosphorylation of either protein following exposure to CM . These findings suggest that, under the conditions tested, the early cellular effects of CM on d-SH-SY5Y cells are not mediated via alterations in pTau (S396) or pIRS1 (S616) signaling (**Supp. Fig. 2**). It is possible that longer exposure to inflammatory stimuli may be required to induce detectable changes in these markers.

### FGA139 prevents LPS-induced NO release in RAW264.7 cells. HMC3 cells do not respond to LPS stimulation

Nitric oxide (NO) is a critical mediator in inflammatory pathways, playing a key role in cellular inflammation and oxidative stress (Cinelli et al., 2020). To explore whether changes in NO secretion from LPS-activated RAW264.7 and HMC3 cells correlate with dendrite morphology alterations in d-SH-SY5Y cells, we evaluated NO levels in CM^+^ from these cell lines, with and without protease inhibitor FGA139.

In RAW264.7 cells, one-way ANOVA showed a significant effect of treatment, (F (2,10) = 182.5; *p* < 0.0001). Further Tukey’s post hoc tests confirmed that LPS stimulation (1 μg/ml, 24h), led to a significant increase in extracellular NO levels from - 3.01 ± 0.06 µM to 14.60 ± 0.58 µM, ([NM vs LPS], p < 0.0001) and that pre-incubation with FGA139 (1 µM) effectively counteracted the LPS effect (10.44 ± 0.87 µM, [LPS vs 1 µM FGA139 + LPS], p = 0.0035) (**Fig. 4A**). On the contrary, LPS incubation did not alter NO secretion from HMC3 cells (**Fig. 4B**).

**Figure 4.**
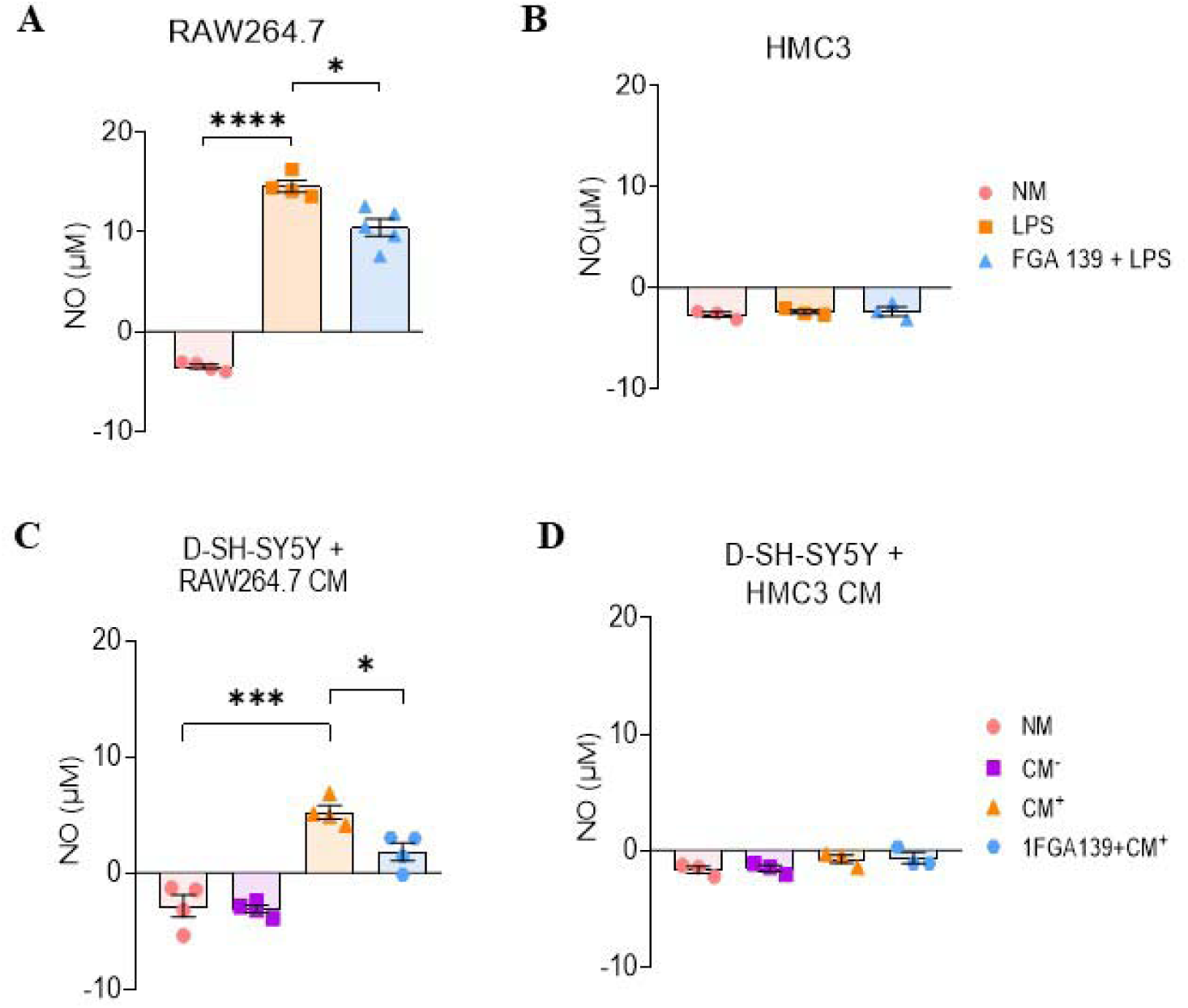
FGA139 Inhibits LPS-Induced Nitric Oxide (NO) Production in RAW264.7 Cells and in d-SH-SY5y cells incubated with RAW264.7 CM. (**A**) NO quantification in RAW264.7 culture media. (**B**) NO levels in HMC3 culture media. (**C**) NO measurement in d-SH-SY5Y cells treated with conditioned media (CM) from RAW264.7. (**D**) NO levels in d-SH-SY5Y cells incubated with HMC3 CM. Experimental conditions: NM (normal media), LPS (1 μM LPS for 24 h in RAW264.7 / 48 h in HMC3) and FGA139 (1 μM ABA for 24 h). Data are presented as mean ± SEM (n = 4-5 per condition) and analyzed by one-way ANOVA followed by Tukey’s multiple comparisons test (**p < 0.01; ****p < 0.0001).

### Macrophage-derived CM induces NO secretion in d-SH-SY5Y cells, attenuated by FGA139

Given that neurons can also secrete nitric oxide (NO) in response to environmental stimuli, we next examined NO levels in d-SH-SY5Y cells following incubation with CM . One-way ANOVA revealed a significant treatment effect after exposure to CM from RAW264.7 cells (F (2,9) = 27.24; p = 0.0002). Tukey’s post hoc analysis showed that 24-hour incubation with RAW264.7 CM significantly increased extracellular NO levels in d-SH-SY5Y cells (from -2.81 ± 0.95 µM to 5.26 ± 0.57 µM; [NM vs CM], p < 0.0001), whereas CM had no effect (−3.07 ± 0.31 µM). Pre-treatment with 1 µM FGA139 significantly counteracted the CM -induced NO elevation (1.83 ± 0.75 µM; [CM vs 1 µM FGA139 + CM], p = 0.0296) (**Fig. 4C**), suggesting that the NO detected in the medium was secreted by d-SH-SY5Y cells in response to macrophage-derived inflammatory signals. In contrast, CM from HMC3 microglial cells after 48 hours did not significantly affect NO secretion in d-SH-SY5Y cells (**Fig. 4D**).

These findings suggest that LPS stimulation induces a more pronounced inflammatory response in macrophages than in microglia, leading to a stronger effect of RAW264.7 CM on neuronal NO production. Furthermore, the ability of FGA139 to attenuate this response underscores its potential as a modulator of macrophage-driven neuroinflammation. Notably, since HMC3 CM also impairs dendritic morphology without increasing NO, alternative mechanisms are likely involved in microglia-mediated neurotoxicity.

### Protease Inhibitor FGA139 Mitigates Oxidative Stress in d-SH-SY5Y Cells induced by Conditioned Media from LPS-Stimulated RAW264.7 Cells

Oxidative stress, characterized by an imbalance between reactive oxygen species (ROS) production and antioxidant defenses, is a key contributor to chronic inflammation (Hussain et al., 2016). To determine whether cysteine protease inhibition confers protection against oxidative stress, we assessed intracellular ROS levels in LPS-stimulated RAW264.7 and HMC3 cells, as well as in d-SH-SY5Y cells exposed to CM from these immune cell lines.

No significant changes in intracellular ROS levels were observed in RAW264.7 (**Fig**. **5A**) or HMC3 cells (**Fig. 5B**) following LPS stimulation or subsequent FGA139 treatment, as determined by one-way ANOVA. However, d-SH-SY5Y cells exposed to CM from LPS-stimulated RAW264.7 cells exhibited a significant increase in ROS levels. One-way ANOVA revealed a significant treatment effect (F (2,6) = 6.206; p = 0.0346), and Tukey’s post hoc test showed that pre-treatment with 1 µM FGA139 significantly reduced ROS levels from 2.78 ± 0.6 µM to 0.15 ± 0.28 µM ([CM vs 1 µM FGA139 + CM], p = 0.0341), even below control levels (NM, 1.35 ± 0.66 µM) (**Fig. 5C**). These results suggest a protective role for FGA139 against ROS-mediated neuronal damage.

**Figure 5.**
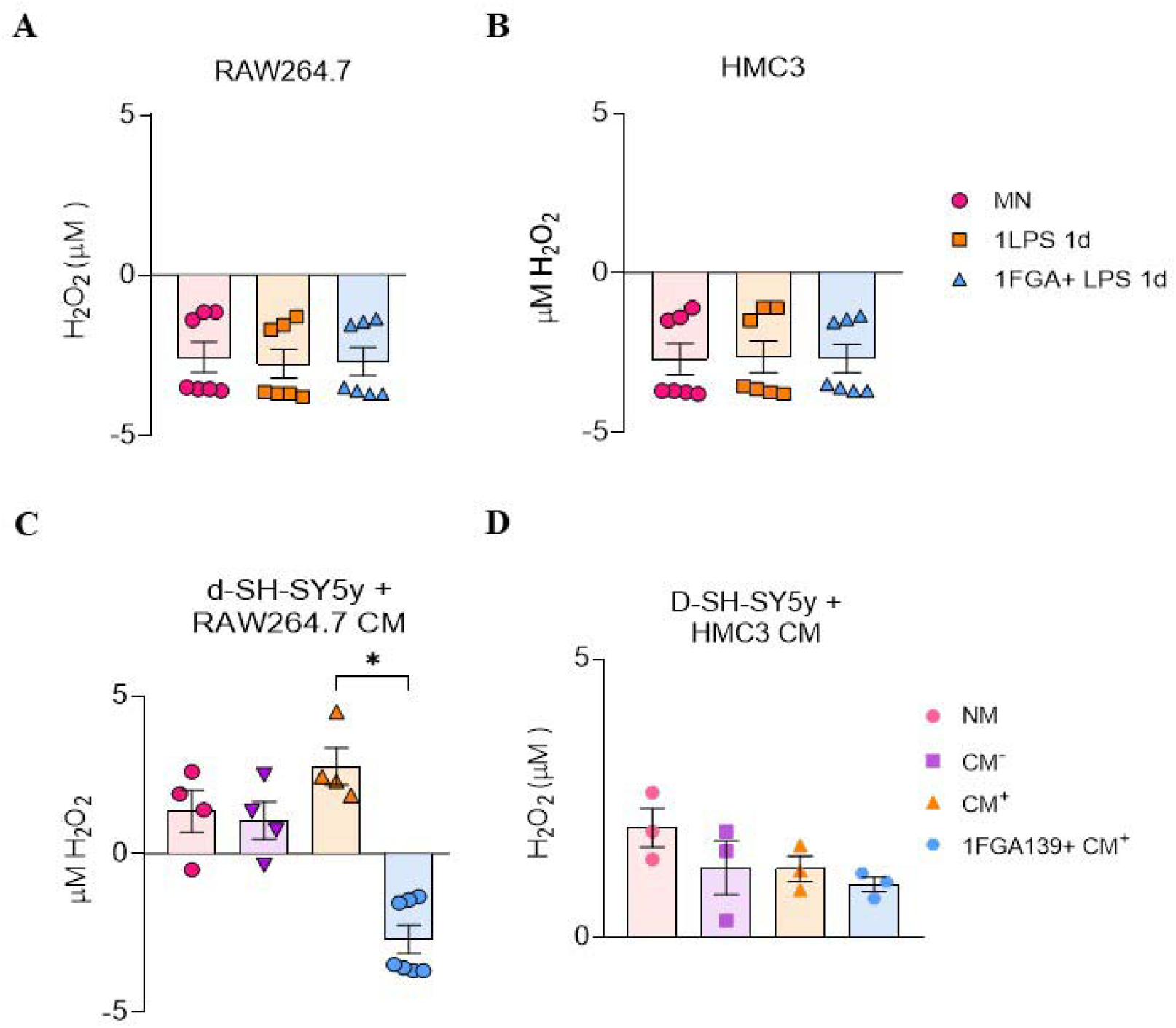
FGA139 Inhibits LPS-Induced Reactive Oxygen Species (ROS) production in d-SH-SY5y cells incubated with RAW264.7 CM. (**A**) Quantification of H O (ROS) levels in RAW264.7 and (**B**) HMC3 cells after 24 h incubation with or without FGA139, followed by LPS treatment for 24 h (RAW264.7) or 48 h (HMC3). (**C**) ROS levels in d-SH-SY5Y cells after incubation with conditioned media (CM) from RAW264.7 and (**D**) HMC3 for 24 or 48 h, respectively, after FGA139 pre-treatment. Data are presented as mean ± SEM (n = 3-7 per condition) and analyzed by one-way ANOVA followed by Tukey’s multiple comparisons test (*p < 0.05).

In contrast, exposure of d-SH-SY5Y cells to CM derived from LPS-stimulated HMC3 cells, with or without FGA139, did not significantly alter ROS levels (**Fig. 5D**).

Collectively, these findings indicate that CM from LPS-activated macrophages (RAW264.7), but not microglia (HMC3), induces oxidative stress in d-SH-SY5Y cells, implying that the neurotoxic mediators are cell-type specific rather than species-specific. The ability of FGA139 to neutralize this oxidative response underscores its potential as a therapeutic modulator in neuroinflammatory conditions.

### FGA139 counteracts TNF-***α*** secretion in LPS-stimulated HMC3 cells, but not in d-SH-SY5Y exposed to CM^+^

To better understand the specific components of CM from HMC3 cells that impact d-SH-SY5Y dendrite length and viability, we assessed levels of tumor necrosis factor-alpha (TNF-α), a well-characterized pro-inflammatory cytokine. TNF-α plays a critical role in immune regulation and is a key pathological mediator in neuroinflammatory disorders such as Alzheimer’s disease and Parkinson’s disease(Subedi et al., 2020). Although microglia are the primary source of TNF-α in the brain, astrocytes and neurons also express and secrete TNF-α, contributing to the inflammatory cascades in neurological disorders (Olmos & Lladó, 2014).

RAW264.7 CM^+^ induced a significant impact on d-SH-SY5y cells TNF-α secretion (one way ANOVA, F (3,8) = 74.57; *p* < 0.0001). Tukey’s post hoc tests confirmed that CM^+^ increased significantly TNF-α extracellular levels (from 1.1 ± 0.52 pg/ml to 44.72 ± 5.66 pg/ml, [NM vs CM^+^], *p* < 0.0001), but FGA139 failed to counteract this effect (**Fig. 6A**). Similarly, HMC3 CM^+^ had a significant impact on d-SH-SY5y cells TNF-α secretion (one Way ANOVA, F (2,6) = 5.91; *p* = 0.0382). Tukey’s post hoc tests revealed that HCM3 CM^+^ led to a significant increase in d-SH-SY5Y extracellular TNF-α levels (from 0.64 ± 0.43 pg/ml to 16.96 ± 4.42 pg/ml, [NM vs CM^+^], *p* = 0.0466), but again, FGA139 pre-treatment did not counteract this increase (**Fig. 6B**).

**Figure 6.**
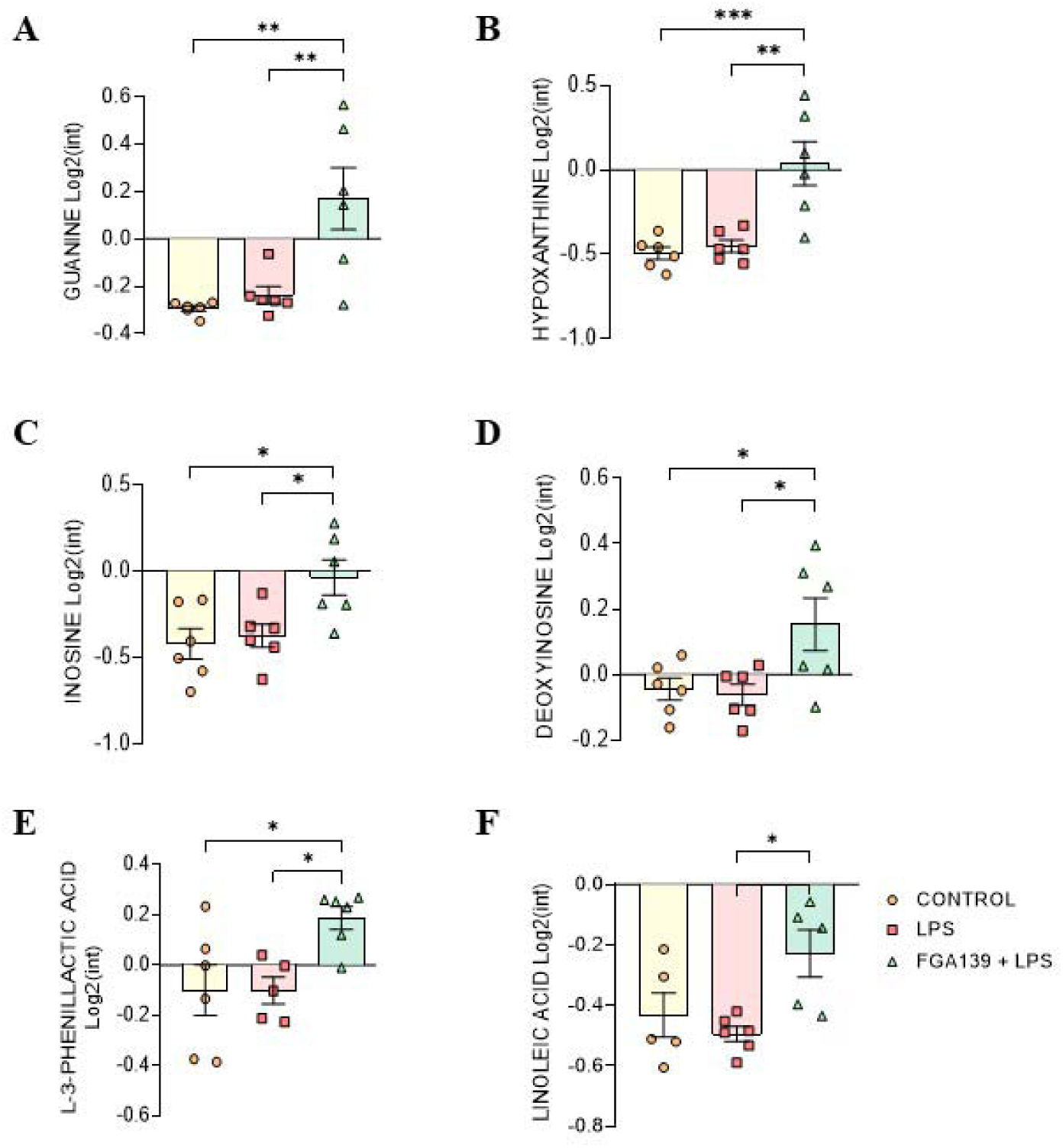
FGA139 does not reduce SHSY5y secreted TNF-U in response to conditioned media, but exhibits a Modest Effect on LPS-Induced TNF-U Secretion in HMC3 Cells. (**A**) TNF-a levels in d-SH-SY5Y cells incubated with conditioned media (CM) from RAW264.7. (**B**) TNF-a levels in d-SH-SY5Y cells incubated with CM from HMC3. (**C**) TNF-a secretion by HMC3 cells. Data are presented as mean ± SEM (n =3 per condition) and analyzed by one-way ANOVA followed by Tukey’s multiple comparisons test (* p < 0.05; **p < 0.01; ****p < 0.0001) and unpaired Student’s t-test (#p < 0.05).

LPS also induced TNF-α secretion in HCM3 cells (one-way ANOVA, F (2,6) = 8.83; *p* = 0.016). Post hoc tests indicated that TNF-α levels increased significantly from 0.061 ± 0.32 pg/ml to 1.91 ± 0.41 pg/mL ([NM vs LPS], *p* = 0.0135). Interestingly, this effect was partially counteracted by pre-treatment with 1 µM FGA139 (0.89 ± 0.14 pg/ml), as these levels were no different from control. A direct t-test comparing [LPS vs 1 µM FGA139 + LPS] confirmed a significant reduction (*p* = 0.0392) (**Fig. 6C**).

These findings indicate that, i) although less responsive than macrophages, HMC3 microglia do secrete TNF-α upon LPS stimulation, and this response can be modulated by FGA139; ii) CM^+^ from both human microglia and murine macrophages induces a robust TNF-α response in d-SH-SY5Y cells, with macrophage CM eliciting a more pronounced effect; iii) TNF-α detected in d-SH-SY5Y cells is of human origin, confirming secretion by the neuronal cells themselves.

Together, these results highlight the anti-inflammatory potential of FGA139 in microglia. However, they also suggest that the neuroprotective effects observed in d-SH-SY5Y cells occur via TNF-α–independent mechanisms.

### Protease inhibitor FGA139 alters the extracellular metabolome in stimulated HMC3 cells unveiling a novel mechanism of action

To further investigate the underlying mechanisms and the potential effect of HCM3 CM^+^ on d-SHSY5y we performed an unbiased metabolomic analysis of the extracellular media from stimulated HCM3, in the presence or absence of protease inhibitor FGA139, at the analytical unit *of IIS La Fe*.

Metabolome analysis could not detect polar metabolite alterations induced by LPS exposure. However, FGA139 significantly impacted guanine metabolites (one-way ANOVA, F (2,15) = 10.38, *p* = 0.0015). Tukey’s post hoc tests showed significant differences between FGA139 + LPS (0.17 ± 0.13) condition with control (−0.29 ± 0.011) [NM vs 1 µM FGA139 + LPS], *p* = 0.0022 and with LPS condition (−0.24 ± 0.036, [LPS vs 1 µM FGA139 + LPS], *p* = 0.0061 (**Fig. 7A**). The treatment had also a significant effect on hypoxanthine levels (one-way ANOVA F (2,15) = 13.44, *p* = 0.0005). Tukey’s post hoc tests showed that pre-treatment with FGA139 significantly increased hypoxanthine levels (0.038 ± 0.13) compared with control (−0.49 ± 0.037, [NM vs 1 µM FGA139 + LPS], *p* = 0.0008) and compared with LPS-treated group (−0.45 ± 0.036, [LPS vs 1 µM FGA139 + LPS], *p* = 0.0017) (**Fig. 7B**). Likewise, the treatment had an effect on inosine levels (one way ANOVA, F (2,15) = 5.87; *p* = 0.0131). Tukey’s post hoc tests showed that cells pre-treated with FGA139 and exposed to LPS significantly increased inosine levels (−0.04 ± 0.101) compared with control (−0.42 ± 0.088, [NM vs 1 µM FGA139 + LPS], *p* = 0.0174) and LPS-treated groups (−0.37 ± 0.067, [LPS vs 1 µM FGA139 + LPS], *p* = 0.0369) (**Fig. 7C**). Deoxyinosine followed the same pattern, (F (2,15) = 4.981; *p* = 0.0219). Tukey’s post hoc tests showed that pre-treatment with FGA139 significantly increased deoxyinosine levels (0.15 ± 0.081) compared with control (−0.044 ± 0.033, [NM vs 1 µM FGA139 + LPS], *p* = 0.0485) and compared with LPS (0.061 ± 0.032, [LPS vs 1 µM FGA139 + LPS], *p* =0.0315) (**Fig. 7D**). Interestingly, a similar pattern was observed in some lipid metabolites. Microglia extracellular L-3-phenyllactic acid (PLA) was significantly affected by treatment (one-way ANOVA F (2,14) = 5.299; *p* = 0.0193). Tukey’s post hoc tests showed that pre-treatment with FGA139 significantly increased PLA levels (0.19 ± 0.045) compared with control (−0.099 ± 0.1, [NM vs 1 µM FGA139 + LPS], *p* = 0.032) and with LPS-treated group (−0.1007 ± 0.054, [LPS vs 1 µM FGA139 + LPS], *p* = 0.0401) (**Fig. 7E**). Similarly, extracellular linoleic acid (LA), an essential ω-6 polyunsaturated fatty acid was significantly affected by the treatment (one-way ANOVA, F (2,13) = 5.42; *p* = 0.0194). Tukey’s post hoc tests confirmed that pre-treatment with FGA139 significantly increases LA levels (−0.23 ± 0.078) compared with LPS-treated group (−0.495 ± 0.034, [LPS vs 1 µM FGA139 + LPS], *p* = 0.0178) but not with control group (**Fig. 7F**).

**Figure 7.**
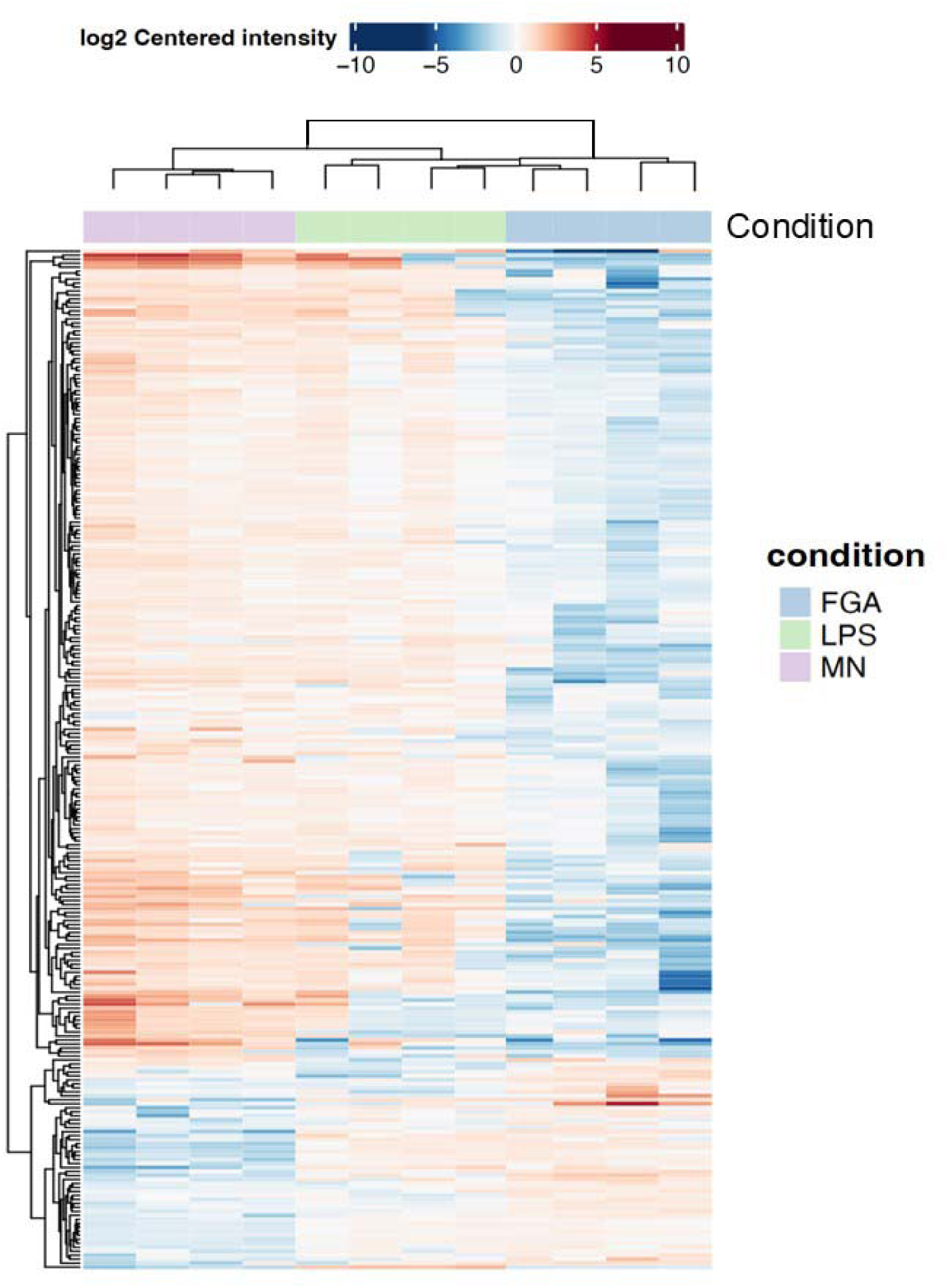
FGA139 Enhances Protective Metabolites in LPS-Stimulated HMC3 Cells. Quantification of guanine (**A**), hypoxanthine (**B**), inosine (**C**), deoxyinosine (**D**), L-3-phenyllactic acid (**E**), and linoleic acid (**F**) in HMC3 cells following FGA139 treatment. Data are presented as mean ± SEM (n = 6 per condition) and analyzed by one-way ANOVA followed by Tukey’s multiple comparisons test (*p < 0.05; **p < 0.01; ***p < 0.001).

These results confirm that HCM3 has a limited response to LPS exposure, with no significant metabolome changes in the extracellular media. On the other hand, our metabolome analysis discloses a novel mechanism underlying cysteine protease inhibitor protective effects via increasing extracellular purine and lipid metabolites with potentially anti-inflammatory effects.

### Untargeted metabolome analysis of HCM3 cells exposed to LPS and FGA139 treatment

In addition to polar metabolome, we performed an untargeted analysis to further characterize microglia response to LPS. The volcano plot of untargeted metabolites did not show great differences. CEUMassMediator identified thymine (with level 2 confidence) as significantly increased by LPS (**Supp. Fig. 3**). Similarly, we analyzed untargeted metabolome comparing FGA139 pretreatment in LPS-stimulated HCM3. The volcano plot displayed a larger number of significantly altered metabolites, also consistent with the greater effect observed in the polar metabolome analysis. Among the differentially expressed metabolites identified by CEUMassMediator (level 2 confidence) we found that creatine, N-phenylacetylglycine, vitamin B6 metabolite 4-pyridoxic acid could be potentially decreased in FGA139 pretreated samples, whereas penicillin G could be increased in those conditions (**Supp. Fig. 4**).

### LPS induces a pro-inflammatory state in human microglia, dominated by interferon signaling and innate immune genes

Following the metabolomic analysis, we conducted comprehensive proteomic profiling of HMC3 cells exposed to LPS and compared their expression patterns to those of cells pretreated with FGA139 prior to LPS stimulation.

Pre-treatment with FGA139 induced a broad suppression of protein expression compared to both non-stimulated and LPS-stimulated conditions, evident in the clustering heatmap panel in blue colors (**Fig. 8**). Interestingly, many of the proteins in this cluster exhibited limited responsiveness to LPS alone. In contrast, the lower quartile cluster revealed proteins whose expression was lower under basal conditions but upregulated in response to both LPS and FGA139. This pattern suggests that some subsets of proteins may be commonly activated under pro-inflammatory and protease-inhibitory conditions. Altogether, the expression profiles indicate that the anti-inflammatory activity of FGA139 does not seem to act by solely counteracting LPS-induced pathways, but rather through broader protein networks regulatory mechanisms. To better understand variations of individual proteins, between LPS-treated and control samples, a volcano plot was generated (**Fig. 9A**). Notably, TMEM188 was the only protein significantly downregulated (three-fold decrease) by LPS treatment. In contrast, all other significantly modulated proteins were upregulated (**Fig. 9B**). Among these, TSC22D3 (also known as GILZ) displayed the highest fold change (11.55-fold increase), followed by LGALS9 (7.89-fold), BST2 (6.41-fold), and OAS2 (4.82-fold). RIG-I exhibited the lowest fold increase, at approximately twofold. Most of the upregulated proteins formed functionally interconnected networks, with the exception of TMEM188 and TSC22D3, as shown in the protein-protein interaction map constructed using the STRING database (**Fig. 9C**).

**Figure 8:**
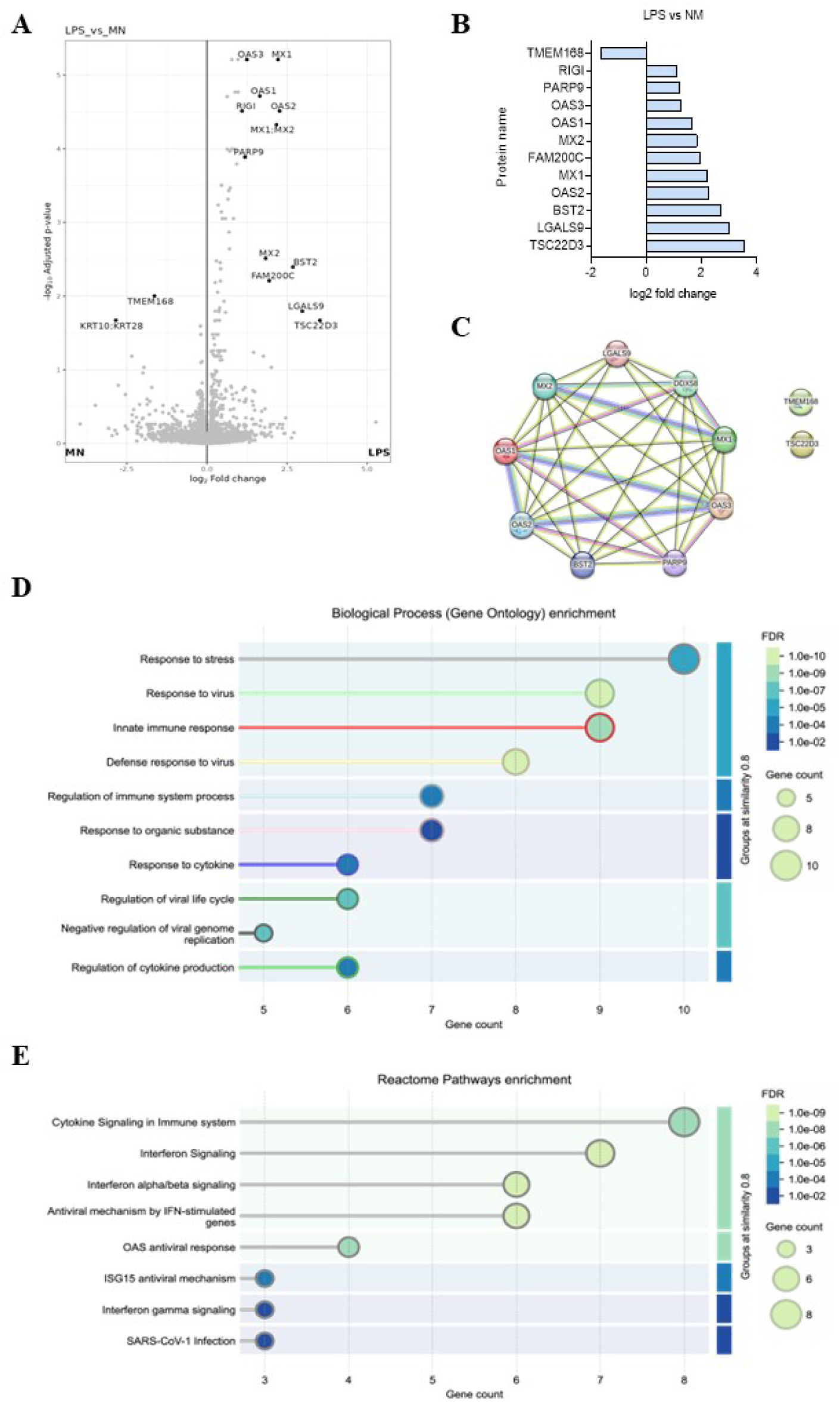
Differential expression analysis. Heatmap showing hierarchical clustering of gene expression profiles across three experimental conditions: MN (purple), LPS (green) and FGA (blue), n = 4 per condition. Rows represent individual genes and columns represent biological samples. Gene expression values represented as log -centered intensities, with red indicating upregulation and blue indicating downregulation relative to the mean expression across samples. Both genes and samples were clustered using hierarchical clustering based on Euclidean distance.

**Figure 9:**
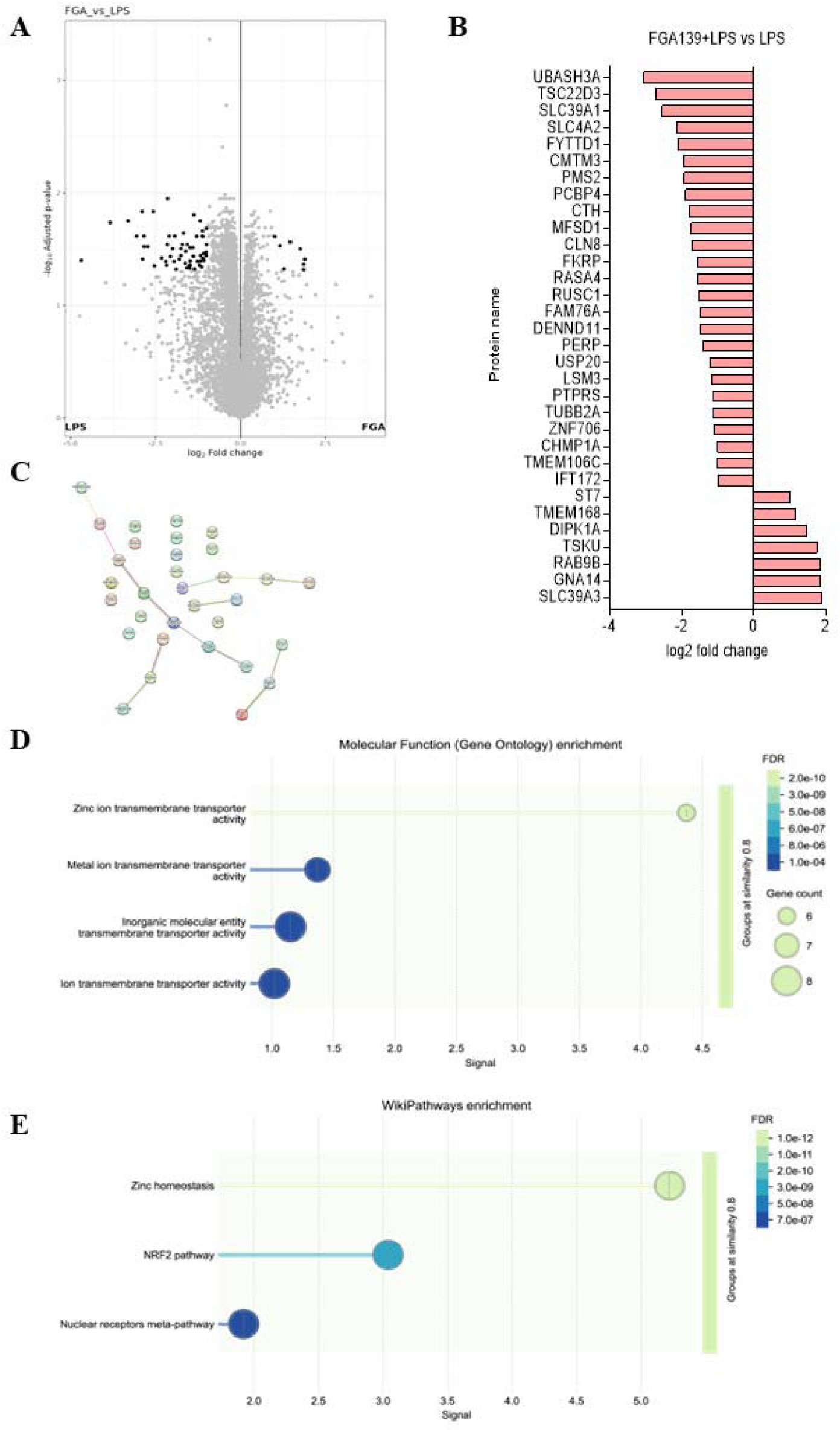
Differential gene expression and functional enrichment analyses comparing NM and LPS conditions. (**A**) Volcano plot showing differentially expressed genes between NM and LPS conditions. The x-axis represents the log fold change, and the y-axis represents the -log adjusted p-value. Highlighted genes show significant changes with strong statistical support. (**B**) Bar plot of the significant genes that came of significant when comparing NM vs. LPS, ranked by log fold change. (**C**) Protein-protein interaction network of significantly different genes constructed using STRING database, illustrating potential functional interactions among key differentially expressed proteins. The lines connecting proteins indicate different types of interactions: Pink, experimentally validated*; L*ight blue, curated database*; D*ark green, gene neighborhood; Red, gene fusions; Dark blue, gene co-occurrence; Light green, text mining; Black, co-expression; Purple, protein homology. (**D**) Gene Ontology (GO) enrichment analysis of biological processes associated with significantly different genes. Bubble size reflects gene count, color scale indicates false discovery rate (FDR), and the x-axis indicates the number of genes per term. (**E**) Reactome pathway enrichment analysis of significantly different genes. Bubble size reflects gene count, color scale indicates false discovery rate (FDR), and the x-axis indicates the number of genes per term. When STRING evaluates predicted protein-protein interactions or enrichments (like GO terms, pathways, or domains), it calculates p-values for each result. Since there are often many comparisons, it adjusts these p-values using FDR correction methods (like Benjamini-Hochberg) to control the chance of false positives. An FDR threshold (like 0.05 or 0.01) means that you accept a 5% or 1% expected proportion of false positives among the results you consider significant. Analysis done with n = 4 per condition.

Biological process enrichment analysis revealed that the differentially expressed proteins were predominantly involved in stress responses and innate immune pathways, consistent with expected LPS-induced effects. Gene Ontology enrichment showed activation of stress-related proteins (e.g., TSC22D3, LGALS9) and antiviral/innate immune effectors (e.g., RIG-I, OAS1/2/3, MX1/2, BST2, PARP9) following LPS stimulation **(Fig. 9D**). Complementarily, Reactome pathway analysis indicated enrichment in cytokine signaling (LGALS9, TSC22D3) and interferon-related pathways (RIG-I, OAS family, MX proteins, BST2, PARP9) (**Fig. 9E**). A full summary of the implicated pathways is presented in **Supp. Table 1**.

Gene ontology enrichment analysis (**Supp. Fig. 5**) comparing LPS-treated to non-stimulated controls revealed strong activation of pathways related to STAT signaling (central to cytokine responses) as well as RNA processing and ATP-dependent activities. These molecular changes are consistent with LPS’s role as a potent activator of innate immunity, initiating broad inflammatory signaling. The enrichment of RNA binding and processing functions suggests enhanced transcriptional and post-transcriptional regulation in microglia during LPS-induced inflammation.

Enrichment in metabolic and mitochondrial components further indicates that LPS may trigger metabolic reprogramming and mitochondrial stress responses. Enrichments related to membrane components agree with LPS-driven remodeling of cellular trafficking, signaling platforms, and organelles.

Notably, LPS stimulation also activated type I interferon signaling and antiviral defense pathways. While LPS is a bacterial ligand, its ability to trigger antiviral programs highlights the extent of the innate immune response, likely mediated through shared pattern recognition receptors and convergent downstream signaling cascades.

Analysis of the proteomic profile of cells pre-treated with FGA139 prior to LPS stimulation (FGA139 + LPS) versus LPS alone revealed that FGA139 predominantly suppresses protein expression, as illustrated by the volcano plot (**Fig. 10A**). This suggests a critical role for cysteine proteases in maintaining essential cell signaling pathways. The most significantly downregulated proteins included UBASH3A (8.3-fold), TSC22D3 (6.6-fold), SLC39A1 (5.9-fold), and SLC4A2 (4.4-fold), whereas the most upregulated were SLC39A3 (3.7-fold), GNA14 (3.6-fold), and RAB9B (3.6-fold) (**Fig. 10B**).

**Figure 10:** Differential gene expression and functional enrichment analysis comparing FGA139 + LPS vs LPS conditions. (**A**) Volcano plot showing differentially expressed genes between FGA139 + LPS vs LPS. The x-axis indicates the log fold change, and the y-axis shows the –log adjusted p-value. Significantly regulated genes are highlighted. (**B**) Bar plot displaying the top differentially expressed genes (FGA139 + LPS vs. LPS), ranked by log fold change. (**C**) Protein–protein interaction network generated using the STRING database, illustrating functional associations among differentially expressed proteins. The lines connecting proteins indicate different types of interactions: Pink, experimentally validated; Light blue, curated database*; D*ark green, gene neighborhood; Red, gene fusions; Dark blue, gene co-occurrence; Light green, text mining; Black, co-expression; Purple, protein homology. (**D**) Gene Ontology (GO) enrichment analysis of molecular functions associated with differentially expressed genes. Bubble size represents gene count, color indicates the false discovery rate (FDR), and the x-axis shows the enrichment signal strength. (**E**) WikiPathways enrichment analysis, highlighting significant pathways such as zinc homeostasis and NRF2 signaling. Bubble size and color represent gene count and FDR, respectively. When STRING evaluates predicted protein-protein interactions or enrichments (like GO terms, pathways, or domains), it calculates p-values for each result. Since there are often many comparisons, it adjusts these p-values using FDR correction methods (like Benjamini-Hochberg) to control the chance of false positives. An FDR threshold (like 0.05 or 0.01) means that you accept a 5% or 1% expected proportion of false positives among the results you consider significant. Analysis done with n = 4 per condition.

Unlike the tightly interconnected proteomic changes triggered by LPS alone, FGA139 pre-treatment led to alterations in protein expression that showed minimal interconnectivity, except for three small clusters of functionally related proteins (**Fig. 10C**). This fragmented pattern may reflect the broad substrate specificity and diverse cellular roles of cysteine proteases, whose inhibition can impact multiple, seemingly unrelated pathways (**Supp. Table 2**).

Gene Ontology (molecular function) enrichment analysis using STRING revealed a pronounced enrichment in proteins associated with zinc transporter activity (**Fig. 10D**). Reactome pathway analysis further indicated that FGA139 affects proteins involved in zinc homeostasis and NRF2 signaling, both known to modulate inflammation (Jarosz et al., 2017) (**Fig. 10E**). Notably, TSC22D3, a gene strongly induced by LPS, was markedly downregulated by FGA139 pre-treatment, while TMEM168, a transmembrane protein suppressed by LPS, was upregulated in FGA139 pretreated microglia.

The overall proteomic profile, including enrichment in terms such as “defense response to virus” and “cytokine-mediated signaling pathway,” suggests that FGA139 redirects microglial responses toward protective defense mechanisms. Backing this, GO enrichment analysis obtained through FragPipe suggests that FGA139, may be involved in attenuating excessive inflammatory cascades while maintaining immune competence through selective modulation of interferon-related pathways. Additionally, the observed upregulation of proteins associated with membrane-bound structures reflect changes in intracellular trafficking, signaling, or structural organization. (**Supp. Fig 6**) Collectively, these findings indicate that cysteine protease inhibition by FGA139 confers a broad protective effect, mitigating LPS-induced alterations in inflammation, transport, and redox regulation, and contributing to the restoration of immune homeostasis under inflammatory stress.

## DISCUSION

Neuroinflammation represents a central mechanism in the pathogenesis of various neurodegenerative disorders. In this study, we investigated the anti-inflammatory and neuroprotective potential of cysteine protease inhibitors, with a focus on FGA139, using *in vitro* models of immune cell-mediated neurotoxicity. Among the tested inhibitors, FGA139 exhibited the most favorable safety profile, validating its selection for further mechanistic studies, in line with prior findings (Agost-Beltrán et al., 2024).

Our results revealed distinct inflammatory responses between RAW264.7 macrophages and HMC3 microglia, which reflect their different origin and function.

A notable observation was the differential sensitivity of RAW264.7 and HMC3 cells to LPS. HMC3 required prolonged exposure to exhibit cytotoxicity, consistent with literature reporting reduced LPS-responsiveness in microglia compared to macrophages (Kocanci et al., 2024). This difference could be attributed to variations in TLR4 expression and downstream signaling pathways (Yi et al., 2020), and would agree with previous reports showing that TLR4 expression in microglia can be induced under ischemic or hypoxic conditions (Yao et al., 2013). Microglia have been shown to respond more robustly to combined LPS + IFN-γ than to LPS alone (Papageorgiou et al., 2016). Despite their limited response, HMC3 cells displayed a significant, though modest, TNF-α release upon LPS stimulation, confirming the presence of functional LPS-responsive receptors (Goksu et al., 2024). Importantly, this effect was attenuated by FGA139, confirming its anti-inflammatory potential.

Likewise, RAW264.7, but not HCM3 cells produced substantial NO in response to LPS. This also agrees with prior reports describing limited inducible NOS, (iNOS) induction in human glial cells by LPS in rodent models (Cinelli et al., 2020; Saha & Pahan, 2006), supporting the idea that macrophages and microglia may engage distinct inflammatory signaling pathways.

Unexpectedly, LPS exposure did not significantly elevate oxidative stress in either RAW264.7 or HMC3 cells, diverging from previous findings (Duque et al., 2025). This discrepancy may result from differences in assay sensitivity, as our colorimetric method lacks the precision of fluorescent probes. Also, other studies report minimal ROS increases in HMC3, unless exposed to combined cytokine stimuli (Dello Russo et al., 2018).

A central aspect of this study involved assessing the impact of conditioned media from LPS-activated immune cells (CM^+^) on differentiated SH-SY5Y cells. Notably, CM from HMC3 but not from RAW264.7 induced a mild reduction in neuronal viability. These intriguing results may reflect different neurotoxic profiles of these two immune cell lines, although we cannot rule out that longer exposure of macrophages CM^+^ can also induce neurotoxicity. Particularly since CM from both immune cell types significantly reduced neurite length in d-SH-SY5Y cells to the same extent, highlighting the ability of inflammatory mediators to induce structural neuronal stress in the absence of overt cytotoxicity. These results are consistent with prior evidence showing that LPS-activated microglia impair neurite integrity (Kaushal et al., 2015; Kocanci et al., 2024). Notably, FGA139 pre-treatment of d-SHSY5Y cells effectively counteracted this neurite shortening, implicating cysteine proteases in inflammation-induced neuronal damage. However, in normal conditions, FGA139 slightly reduced neurite length, suggesting that cysteine proteases also contribute to normal neuronal function and their inhibition is not beneficial in the absence of inflammatory ambience. For instance, extracellular cathepsin L and calpain-1 have been shown to regulate neurite outgrowth and synaptic plasticity (Knopp et al., 2021).

These data suggest that the protective effect of FGA139 likely arises from mitigating overactivation of cysteine proteases, which could amplify inflammatory cascades during stress (Pišlar et al., 2021). Calpain, for example, is upregulated in activated microglia and it is linked to myelin loss, calcium dysregulation, and neuronal death (Vosler et al., 2008). Caspases and cathepsins similarly promote inflammatory damage by processing pro-inflammatory mediators (Campden & Zhang, 2019; Van Opdenbosch & Lamkanfi, 2019). Our data further supports this mechanistic link and emphasizes the potential of inhibiting cysteine proteases as a therapeutic approach (Siklos et al., 2015).

Despite pronounced changes in neurite morphology, exposure to CM did not significantly elevate tau phosphorylation at Ser396 or IRS1 phosphorylation at Ser616, markers associated with neurodegeneration and metabolic stress, respectively (Chou et al., 2020; Gao et al., 2018). This may be attributed to the short exposure period or insufficient concentration of toxic mediators. Prolonged exposure might be necessary to reveal such changes. Alternatively, the SHSY5y differentiation protocol might influence tau isoform expression, and it has been reported that enhanced hyperphosphorylation typically requires RA+BDNF treatment (de Medeiros et al., 2019).

Interestingly, CM from RAW264.7, but not HMC3, increased ROS and NO levels in SH-SY5Y cells. This aligns with the established role of macrophage-derived factors in disrupting neuronal redox balance (Chiu et al, 2021) and suggests a neuronal nitric oxide synthase (nNOS) activation (J. Chen et al., 2004).

FGA139 effectively mitigated this ROS and NO increase and would align with previous reports supporting the role of calpain inhibitors suppressing ROS production (X. Liu et al., 2022; Zhang et al., 2025). Also, because excessive nNOS activity can exacerbate neuronal damage (L. Zhou & Zhu, 2009), the ability of FGA139 to prevent this NO rise underscores its neuroprotective potential.

However, these results do not correlate with viability (affected by HCM3 but not RAW264.7 CM^+^) and dendrite length (that is affected by both). These findings may reflect species-and cell-type-specific differences in secreted inflammatory neurotoxic mediators (Wolfe et al., 2018). Furthermore, microglia-induced neurotoxicity depends on the nature of the activating stimulus, for instance Aβ-stimulated HMC3 cells can induce pronounced neurodegeneration (Wei et al., 2022), likely greater than LPS-stimulated HCM3.

Strikingly, CM exposure from both immune cell lines elevated TNF-α release from SH-SY5Y neurons. Although microglia are considered the primary source of brain TNF-α, it is also expressed and released by astrocytes and neurons (Park & Bowers, 2010). For instance, evidence supports that neurons can express TNF-α, under ischemic conditions (T. Liu et al., 1994), or in Alzheimer models (Janelsins et al., 2008). This neuronal-derived TNF-α may reflect a disrupted microglia-neuron cross talk (Jurgens & Johnson, 2012). It is well known that TNF-α is secreted by stimulated RAW.264.7 cells (Swantek et al., 1999), thus it is plausible that this cytokine could mediate the reduced dendrite length in d-SHSY5Y (Y. Liu et al., 2017).

Furthermore, elevated brain TNF-α facilitate the infiltration of inflammatory cells that can further exacerbate tissue damage ref. Interestingly, RAW264.7-derived CM exhibit greater effect, suggesting that neuronal TNF-α production may be more sensitive to macrophage than macroglia derived damage associated molecular patterns (DAMPs). The consequences would be that the infiltrating macrophages may exponentially exacerbate brain inflammatory milieu.

Intriguingly, FGA139 inhibited microglial TNF-α release, but failed to suppress neuronal TNFα increment, suggesting that the regulation of neuronal TNF-α production may differ from microglia in terms of triggers and control mechanisms, and likely independent of cysteine protease activity. While cysteine proteases have not been reported in the canonical pathway of TNF-α synthesis, our findings support the modulation of cysteine proteases in inflammatory processes in microglia cells, but not neuronal cells.

Metabolomic profiling of HMC3 supernatants revealed no significant changes in polar metabolites following LPS treatment, reflecting the limited LPS responsiveness of this cell line, similar to NO, and ROS. However, pre-treatment with FGA139 prior to LPS exposure induced a significant increase in purine metabolites (guanine, hypoxanthine, inosine, and deoxyinosine). Although astrocytes are considered the primary source of extracellular purines (Ciccarelli et al., 2001), our findings suggest that microglia may be an important contributor to the reparative role associated with extracellular purine metabolites. Also, most purine anti-inflammatory effects have focused on adenosine-based nucleosides, however, our data supports the emerging neuromodulator role of guanine-based purines (Ortiz et al., 2015; Tasca et al., 2018). Elevated hypoxanthine levels have been associated with a reduced Alzheimer’s disease risk (Chouraki et al., 2017), while deoxyinosine exhibits anti-inflammatory effects in preclinical arthritis models, and it is consistently reduced in rheumatoid arthritis patients (Xu et al., 2024). The mechanisms underlying this finding may imply different pathways, like reduced nucleotide degradation, enhanced purine salvage, or increased trafficking. This finding suggests that cysteine proteases may have previously unrecognized roles in nucleotide metabolism, potentially linking proteolytic processes with purinergic signaling, which is important in inflammation and immune responses.

FGA139 also increased extracellular linoleic acid (LA), a polyunsaturated fatty acid with documented anti-inflammatory and remyelination-promoting properties (Whelan & Fritsche, 2013; Zhou et al., 2023). Furthermore, LA derivatives α-dimorphecolic acid (α-DIPA) display neuroprotective effects (Zhu et al., 2025). While the precise role of cysteine proteases in lipid metabolism is unclear, their involvement in lipid droplet turnover and membrane remodeling suggests a plausible link (Ballabio & Bonifacino, 2020). Surprisingly, FGA139 elevated phenyl lactic acid (PLA) levels, a metabolite with antimicrobial and immune-modulating properties (Oezguen et al., 2022). PLA primary source in humans is gut microbiota (Abdul-Rahim et al., 2021), and though PLA’s neuroinflammatory role remains largely uncharacterized, its supplementation has shown beneficial impact in rodent models (Rastogi & Singh, 2022) and structurally related compounds have shown anti-inflammatory and neuroprotective effects (Chen et al., 2011; Kim et al., 2024). To our knowledge, this is the first demonstration of the implication of cysteine proteases in PLA secretion regulation in microglial cells.

Untargeted metabolomic analysis revealed several putatively annotated features (level 2 confidence) that were reduced in FGA139-pretreated LPS-stimulated microglia, compared to LPS alone, those included extracellular levels of N-phenylacetylglycine (PAG), creatine, and 4-Pyridoxic acid (PA). PAG is a glycine conjugate derived from phenylalanine and tyrosine metabolism. If validated, the observed reduction in PAG following FGA139 treatment could suggest a novel anti-inflammatory mechanism, as stress-induced tyrosine catabolism has been linked to mitochondrial toxicity in rodent models (Doessegger et al., 2013) and neuroinflammation (Thau-Zuchman et al., 2022). The role of creatine in neuroinflammation remains debated: while creatine supplementation has shown neuroprotective effects in Parkinson’s disease (PD) models (Leem et al., 2024) and healthy humans (Gutiérrez-Hellín et al., 2025) elevated creatine and related metabolites have also been implicated in PD pathophysiology (Wasilewski et al., 2025). If confirmed, our results could reflect excessive energy demand in activated microglia, attenuated by cysteine protease inhibition via FGA139. Finally, 4-Pyridoxic acid (PA) is a catabolic end-product of vitamin B6 metabolism, because elevated PA has been associated with oxidative stress (Ulvik et al., 2012), the modulation of PA levels by FGA139 could underlie its potential to counteract inflammatory oxidative processes. Future studies using isotopic labeling or chemical standards are needed to confirm these putative annotations and elucidate their functional roles.

Taking together, these findings indicate that FGA139 significantly influences the extracellular metabolic profile of LPS-activated microglia. This shift in metabolic phenotype may contribute to the anti-inflammatory and neuroprotective properties of cysteine protease inhibitors in neuroinflammatory disorders (Lowry & Klegeris, 2018). Proteomic analysis revealed that under control conditions, inflammatory markers were expressed at low levels in HCM3, consistent with a resting, homeostatic microglial phenotype. In contrast, LPS-treated microglia exhibited a marked upregulation of proinflammatory proteins, reflecting a shift toward an M1-like phenotype. Notably, among these, TSC22D3 (GILZ) was significantly induced by LPS. While traditionally associated with anti-inflammatory functions via NF-κB inhibition (Berrebi et al., 2003), emerging evidence suggests a context-dependent role for GILZ. Recent studies in co-culture systems have implicated GILZ as a downstream effector of macrophage migration inhibitory factor (MIF), contributing to both inflammasome activation and M2-like polarization in macrophages (Zhang et al., 2025). Our findings support this role, since GILZ is upregulated in response to LPS but significantly downregulated by FGA139. This suggests that cysteine protease inhibition may attenuate neuroinflammation in part by suppressing GILZ-mediated inflammasome activity. This is a novel mechanistic insight linking protease inhibition to immunomodulation.

Interestingly, pre-treatment with FGA139 selectively modulated the expression of proteins across diverse, seemingly unrelated biological pathways. This finding is consistent with the pleiotropic roles of cysteine proteases in regulating multiple cellular processes. Of particular interest was FGA139 regulation of Zinc transporter family members. FGA139 upregulates SLC39A3 (ZIP3) and downregulates SLC39A1 (ZIP1). This result could be paradoxical; on one hand, ZIP3 can facilitate zinc influx into the cytosol (Jeong & Eide, 2013), and as zinc is a crucial modulator of inflammatory signaling, inhibiting NF-κB and activating PPARγ (Gammoh & Rink, 2017), our findings would imply that FGA139 may exert anti-inflammatory and neuroprotective effects by enhancing cytosolic zinc availability.

On the other hand, SLC39A1 (ZIP1), a broadly expressed plasma membrane zinc importer has been associated with age-related changes in the brain (Olesen et al., 2016). Notably, dysregulation of ZIP1 has been linked to increased susceptibility to neurodegeneration and amyloid-β aggregation (Lang et al., 2012; Walsh et al., 2002). Thus, the suppression of ZIP1 expression by FGA139 could modulate zinc-induced excitotoxicity, adding a further layer of neuroprotective potential mediated through zinc homeostasis. Zinc physiological levels tend to maintain homeostasis and provide anti-inflammatory effects, while excessive zinc during injury or disease can contribute to neurotoxicity through direct and indirect mechanisms, this dual nature makes zinc signaling a complex regulated system. Together, these findings suggest a previously unrecognized role for cysteine proteases in regulating zinc transporter expression (ZIP1, ZIP3), a mechanism potentially critical for the fine tuning of Zinc in neuroinflammatory responses.

Additionally, FGA139 downregulated SLC4A2 (AE2), an electroneutral Cl /HCO exchanger (Ibrahim et al., 2017). AE2 has been implicated in oxidative stress responses and apoptosis under via ROS and caspase-3 activation (Remigante et al., 2022). By downregulating AE2, FGA139 may protect against inflammation-induced oxidative stress and ionic imbalance.

Interestingly, FGA139 upregulated vesicular trafficking proteins like RAB9B, a small GTPase involved in late endosome–Golgi trafficking (Hutagalung & Novick, 2011). It is tempting to speculate that this upregulation could facilitate the vesicular secretion of the diverse small metabolites found in the metabolome analyses of microglial supernatant. The upregulation of TSKU (Tsukushi), a small leucine-rich proteoglycan modulating TGF-β, FGF/MAP kinase, and Wnt signaling pathways (Dellett et al., 2012; Morris et al., 2007; Ohta et al., 2011) would support a trophic role of cysteine protease inhibition under inflammatory conditions.

Also, GNA14 (Gα14) a Gq alpha subunit was also found upregulated in FGA139 treated stimulated microglial cells. Although its role is not well characterized in microglia, Gα14 has been linked to activation of NF-κB signaling (A. M. F. Liu & Wong, 2005), and STAT3 phosphorylation (Lee et al., 2012). Our findings would support a novel role for protease inhibition modulating inflammation via Gq α.

On the other hand, FGA139 pretreatment was found to downregulate proteins implicated in oxidative stress regulation like cystathionine γ-lyase (CTH), a recognized NRF2 target involved in hydrogen sulfide (H S)-mediated antioxidant defense mechanisms (Jamaluddin et al., 2022). Surprisingly, the most significantly downregulated protein was UBASH3A, a negative regulator of NF-κB activity in T cells (Ge et al., 2017), its suppression by FGA139 warrants further investigation, as it may reflect microglia-specific regulatory dynamics during inflammation and protease inhibition.

## CONCLUSION

In summary, this study supports the hypothesis that the novel cysteine protease inhibitor FGA139 exerts anti-inflammatory and neuroprotective effects and prompts for future validation in animal models of disease. In addition, our data confirms important distinctions between microglia and macrophage responses to LPS stimulations and different contributions to neuroinflammation-induced neuronal damage.

Our study reveals unexplored mechanisms of cysteine proteases inhibition in LPS-stimulated microglia, including significantly elevated extracellular purine metabolites and lipids (PLA and LA) with potential anti-inflammatory effects. Proteomic analysis revealed previously unidentified inflammatory mediators including zinc transporters, ion channels, G-protein signaling, vesicular trafficking, and survival pathways. Our interpretations point to potential pathways regulated by cysteine protease inhibition. We acknowledge the limitation of the lack of specificity of the inhibitor, which limit the interpretation of the pathways involved.

Our findings emphasize the therapeutic potential of modulating cysteine proteases to mitigate neuroinflammation. Further investigation of these newly identified metabolite pathways and proteomic profile will enhance understanding of microglial-neuronal interactions contributing to targeted therapies for neuroinflammatory disorders.

## Declaration of generative AI in scientific writing

During the preparation of this work, the authors used AI assistance to revise grammar and correct typographical errors. The authors subsequently reviewed and edited the content as necessary and take full responsibility for the final version of the manuscript.

## Acknowledgments

This work was supported by *Plan Propi* UJI UJI-B2021-21 to AMSP and Universtitat Jaume I, ref GACUJIMC/2024/05 to AMSP. The authors want to thank the donations through the crowdfunding campaign “*Biotecnologia contra el Alzheimer*” organized by UJI (2023). FVG thanks Generalitat Valenciana (PROMETEO with ref. CIPROM/2021/079) for funding. Authors thank Dr. Laura Agost-Beltran for the provision of FGA139 inhibitor. We acknowledge the services of Metabolomics in IIS La Fe, Valencia (Spain) and Proteomics SCSIE University of Valencia, Valencia (Spain), for the analysis and help managing the results.

## Supplementary figures

**Supplementary Figure 1A.**
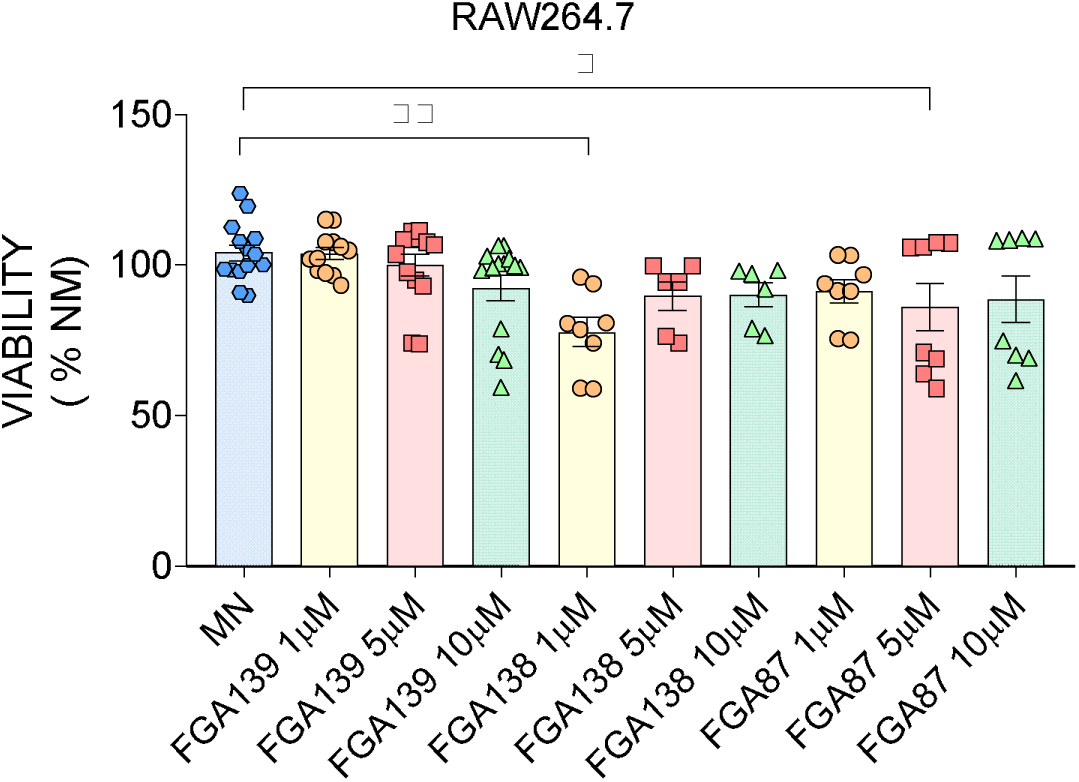
Cysteine proteases inhibitors effect on RAW 264.7 viability. To determine the potential cytotoxicity of our protease inhibitors, 1 µM and 5 µM concentrations of FGA139, FGA87, FGA138 were tested on murine macrophages RAW264.7. The treatment had a significant effect on RAW264.7 cell viability (one-way ANOVA F (4,39) = 5.14; p = 0.002). FGA138 1 µM significantly reduced viability (77,73 ± 4,88 %) compared to control (104,1 ± 2,61 %) (Tukey post hoc test [NM vs 1 µM FGA138], p = 0.0011). Also, FGA87 5µM reduced viability (86,15 ± 7,85%) significantly ([NM vs 5 µM FGA87], p = 0.0445). FGA139 did not affect viability at any concentration tested.

**Supplementary Figure 1B.**
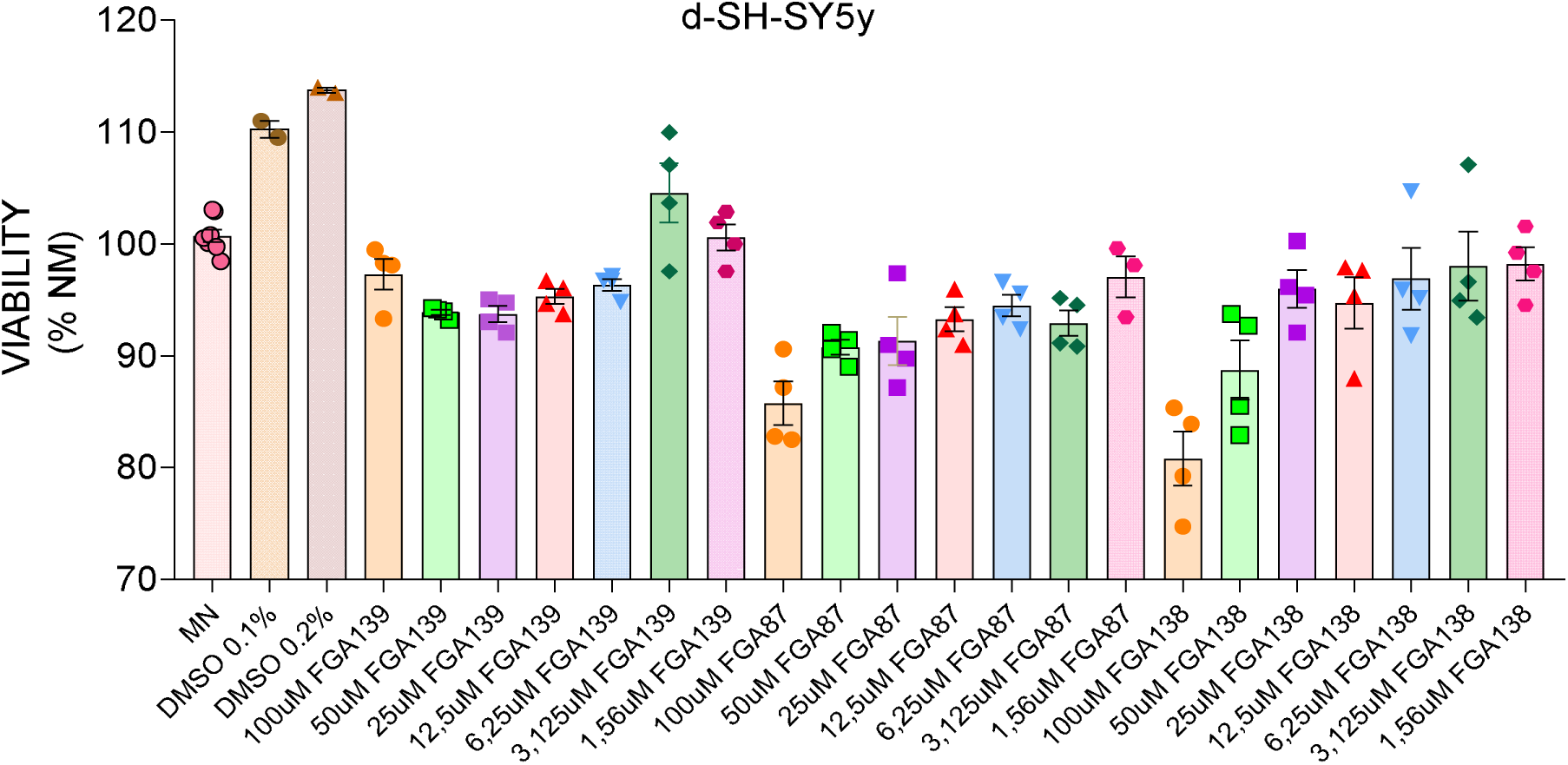
Cysteine proteases inhibitors effect on d-SH-SY5y cells viability. FGA139, FGA138, and FGA87 (ranging from 100 µM to 1.56 µM) were tested on d-SH-SY5Y cells viability. Data are presented as mean ± SEM, analyzed by one-way ANOVA followed by Tukey’s multiple comparisons test (*p < 0.05; ***p < 0.001).

**Supplementary Figure 2.**
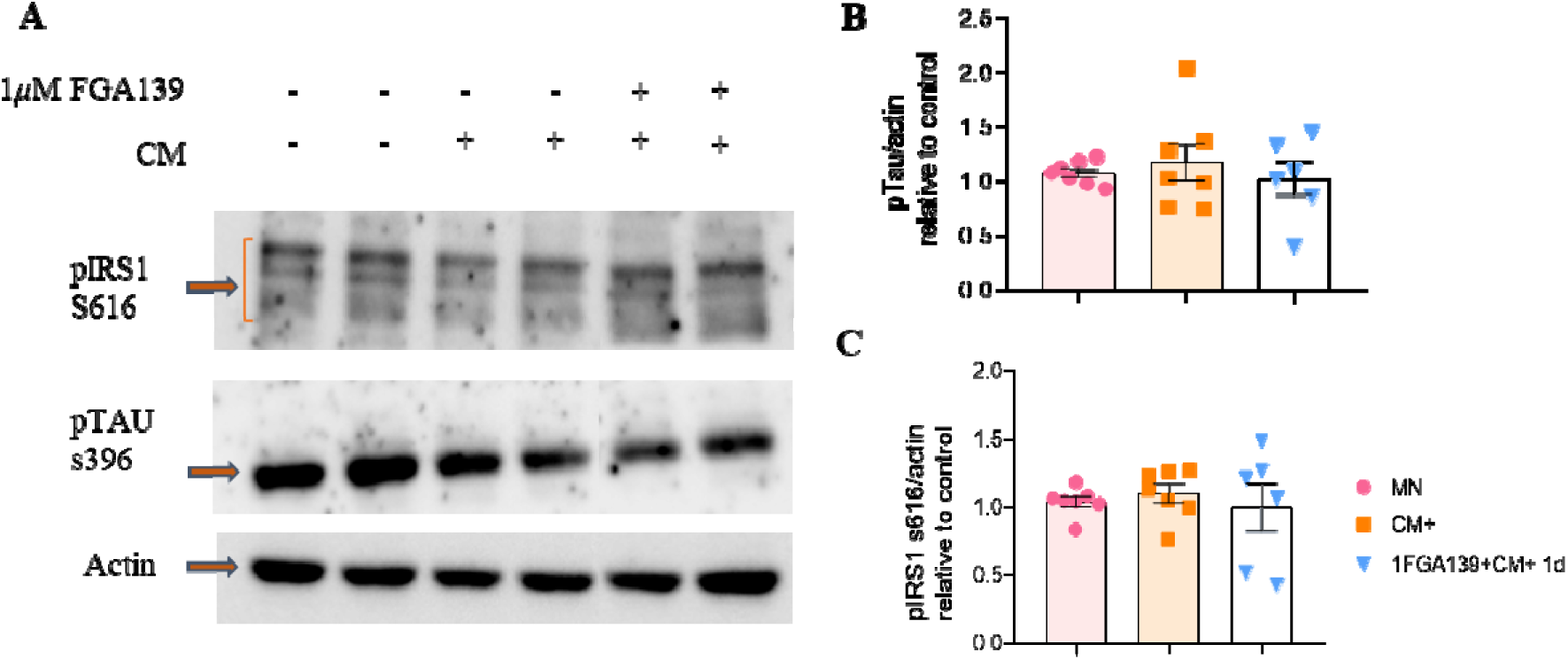
Western Blot Analysis of p-Tau S396, and pIRS1. d-SH-SY5y cells were pre-treated with FGA139, followed by incubation with RAW264.7 CM^+^. (**A**) Representative Western blot images. Actin was used as loading control. (**B**) Quantification of pTAU S396 levels under different conditions as indicated. (**C**) Quantification of pIRS1 S616 levels across tested conditions. Data are presented as mean ± SEM, analyzed by one-way ANOVA.

**Supplementary Figure 3.**
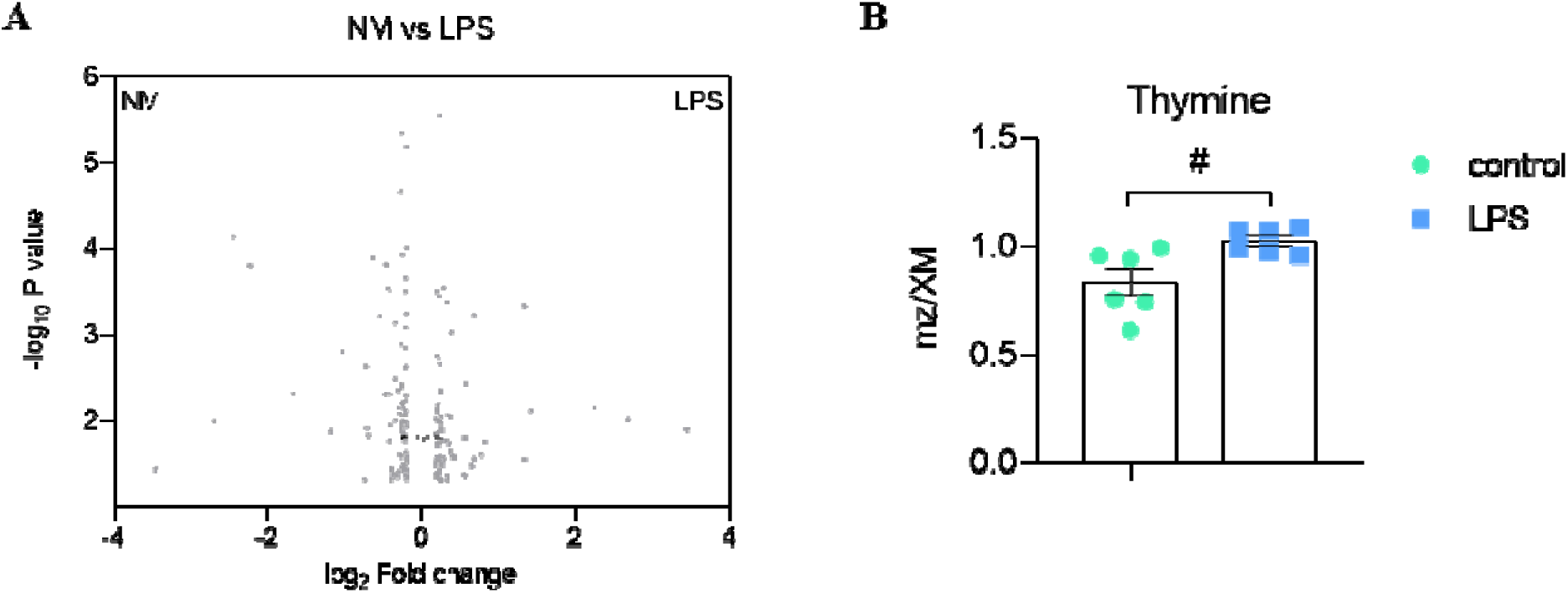
Untargeted metabolomic analysis of microglia supernatant, comparing Normal Media to LPS exposure conditions. (**A**) Volcano plot showing differential metabolite expression between normal medium (NM) and LPS treatment. (**B**) Relative concentrations (mz/XM) in control (0.83 ± 0.061) and LPS treated (1.03 ± 0.023) were significantly different (unpaired Student’s t-test [NM vs LPS], p = 0.0161), indicating that LPS treatment may increase thymine metabolism or secretion.

**Supplementary Figure 4.**
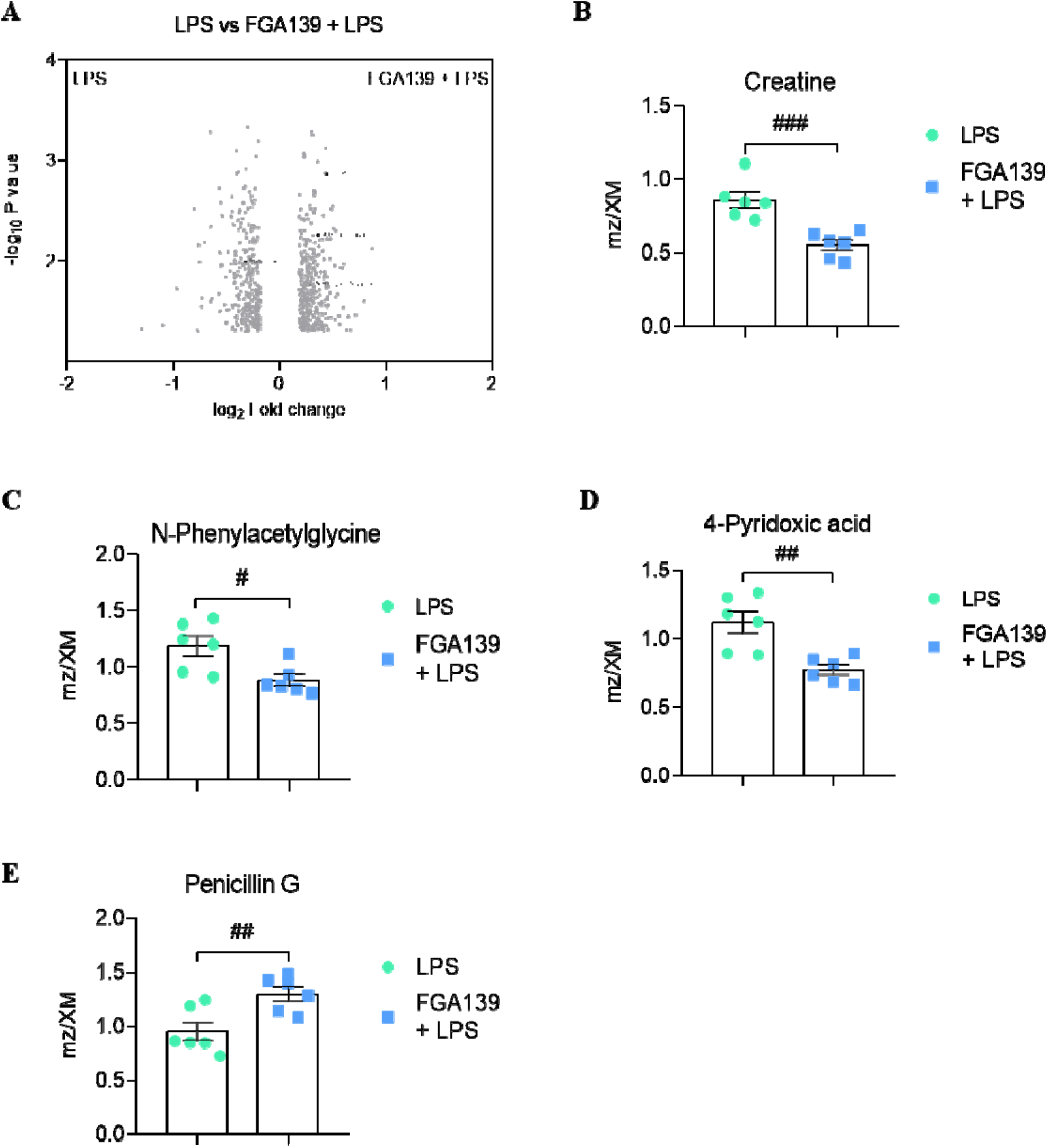
Untargeted metabolomic analysis of microglia supernantant, comparing LPS exposure with and without FGA139 pretreatment. **(A**) The volcano plot comparing these conditions displayed a greater number of significantly altered metabolites, suggesting that FGA139 modifies the LPS-induced metabolic response. (**B**) Among the differentially expressed identified metabolites with level 2 confidence, creatine relative concentration levels (mz/XM) were significantly decreased in FGA139 + LPS (0.56 ± 0.036) compared to LPS alone (0.86 ± 0.055) (Student t-test [LPS vs 1FGA139 + LPS], p = 0.001)**. (C)** Similarly, N-phenylacetylglycine concentrations were markedly reduced when cells were pre-treated with FGA139 (0.088 ± 0.052) compared to LPS alone (1.19 ± 0.087) (Student t-test [LPS vs 1FGA139 + LPS], p = 0.0135). (**D**) The vitamin B6 metabolite 4-pyridoxic acid also exhibited significantly decreased with FGA139 pre-treatment (0.78 ± 0.038) compared to LPS alone (1.12 ± 0.8) (Student t-test [LPS vs 1FGA139 + LPS], p = 0.0029). (**E**) Interestingly, in contrast to the reducing effect observed with other metabolites, penicillin G levels were significantly elevated with the pre-treatment of FGA139 (1.3 ± 0.066) compared to control (0.95 ± 0.086) (Student t-test [LPS vs 1FGA139 + LPS], p = 0.0094. This finding is atypical in this system, as all cultures are supplemented with antibiotics in the culture media.

**Supplementary Figure 5.**
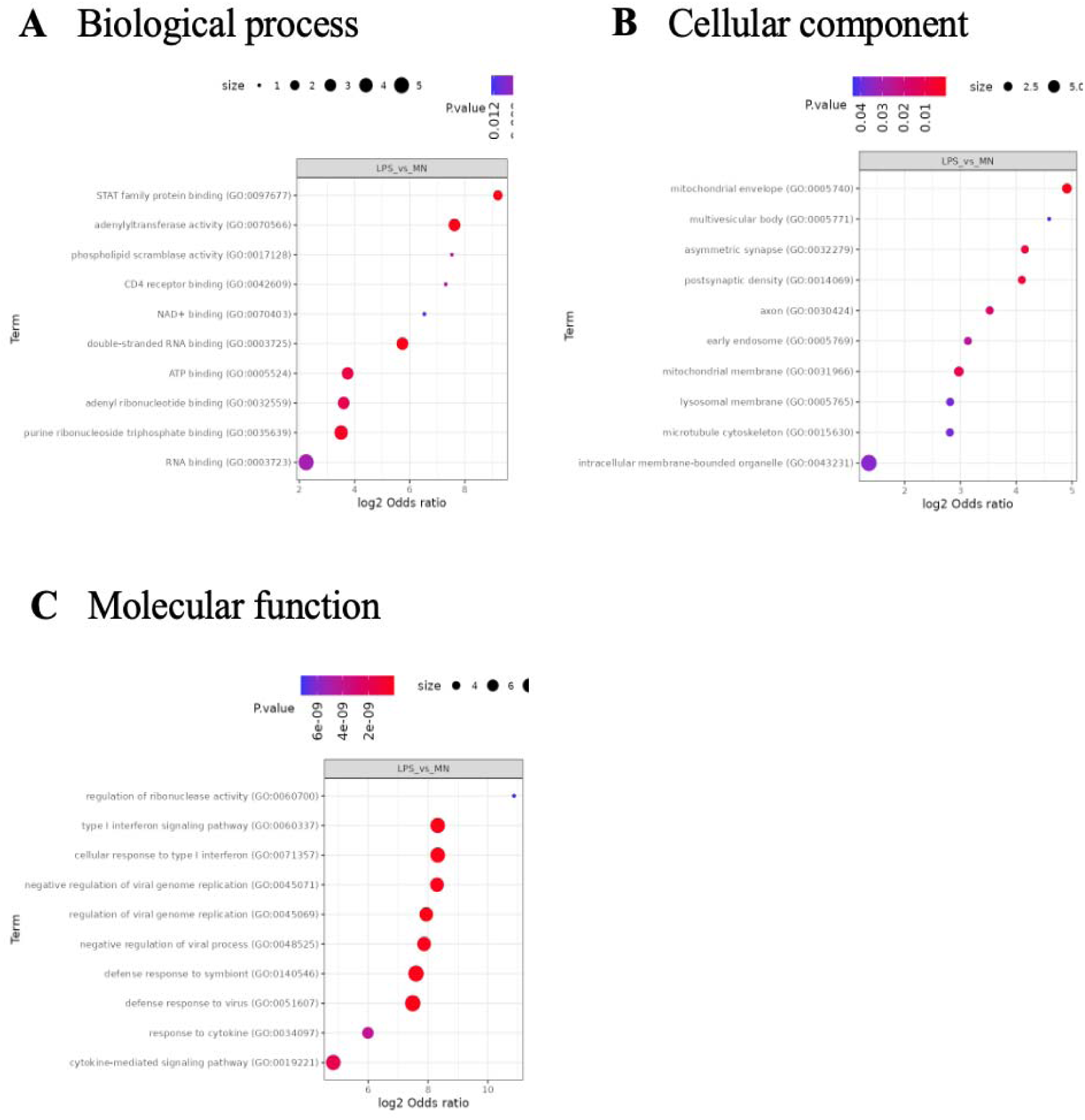
Functional enrichment analysis obtained by FragPipe, comparing LPS vs NM (normal medium) treatment. (**A**) This Gen Ontology (GO) term, *Biological Process* reveals key functional pathways that are more represented, upregulated or activated in the conditions mentioned before. The STAT family protein binding showed the strongest enrichment (logl odds ratio >7, p < 0.01), suggesting a cytokine signaling. This pathway was enriched by adenylyltransferase activity, double-stranded RNA binding, and ATP bindin*g* (logl odds ratios >4, p < 0.01), which suggests active immune signaling, especially innate immunity and cytokine pathways, potential response to a viral mimic, infection, which is coherent with microglia exposed to an inflammatory stimulus. (**B**) Regarding the *cellular compartment* functionally enriched in the LPS condition. Notably, the mitochondrial envelope (GO:0005740) exhibited the highest enrichment (logl odds ratio ∼5), a low p-value, and involved a large number of proteins, indicating substantial mitochondrial remodeling in response to LPS. Similarly, components such as the multivesicular body (GO:0005771) and lysosomal membrane (GO:0005765) were also highly enriched and statistically significant, reflecting increased vesicular trafficking and degradative activity typical of microglial activation. Surprisingly, we found enrichment of synaptic structures, including the asymmetric synapse (GO:0032279) and postsynaptic density (GO:0014069). These findings collectively highlight a robust shift in cellular architecture, particularly within mitochondrial and endolysosomal pathways, accompanying microglial activation by LPS. (**C**) *Molecular functions* upregulated in LPS-treated condition, revealed robust activation of immune-related biological processes. Immune response pathways such as the type I interferon signaling pathway (GO:0060337) and cellular response to type I interferon (GO:0071357) were highly enriched (log OR ∼8), statistically significant (deep red), and involved a large number of proteins, as indicated by the dot size. Additionally, pathways like defense response to virus (GO:0051607), regulation of viral genome replication (GO:0045069), and cytokine-mediated signaling pathway (GO:0019221) also demonstrated strong enrichment and significance. These findings highlight a potent antiviral and pro-inflammatory transcriptional shift in LPS-stimulated microglia, characterized by heightened cytokine responses and activation of interferon-stimulated pathways. *Y-axis: Lists Gene Ontology (GO) terms. X-axis (log2 Odds ratio): Represents the enrichment of each term. Higher values mean the term is more enriched (above than expected by chance). Dot color: reflects p-value significance according to the gradient. Dot size: reflects the number of genes involved (likely the count of genes associated with the term that are enriched)*.

**Supplementary Figure 6.**
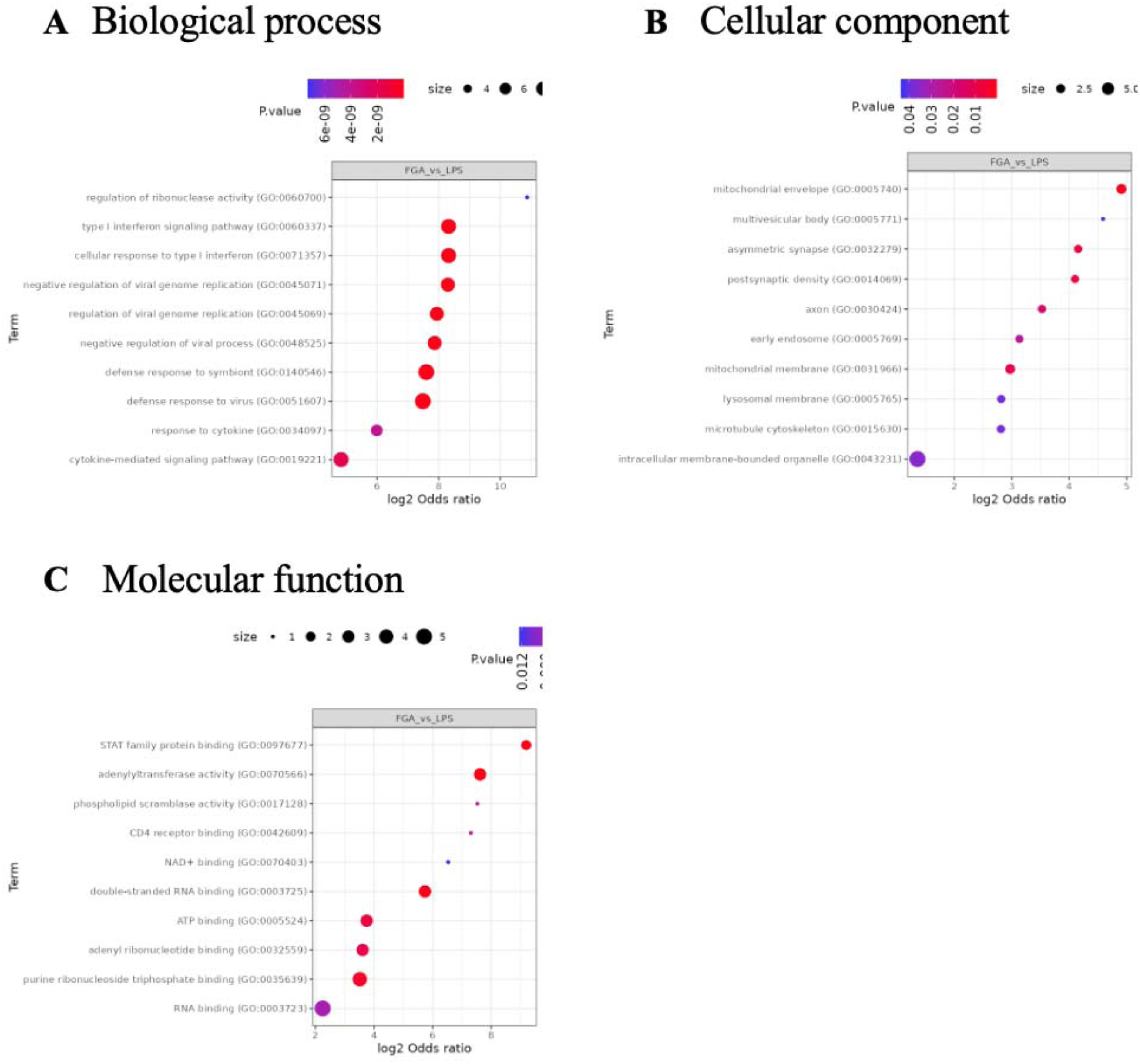
Functional enrichment analysis obtained by FragPipe, comparing FGA139+LPS vs LPS treatment. **(A)** This GO enrichment analysis (Biological Processes) reveals key functional pathways that are more represented, upregulated or activated) in FGA139 pre-treated activated microglia compared to those treated with LPS alone. These represent a distinct shift toward antiviral and regulatory immune responses, diverging from the classical pro-inflammatory profile typically induced by LPS. The upregulation of type I interferon signaling pathways (GO:0060337) and cellular response to type I interferon (GO:0071357), both highly enriched and statistically significant with broad protein involvement, suggests that FGA139 may promote a controlled, antiviral-like state rather than a generalized inflammatory response. Additionally, the marked enrichment of regulation of ribonuclease activity (GO:0060700)—the top hit with the highest logl odds ratio (∼10)—implies a post-transcriptional regulatory mechanism may be at play, possibly helping to fine-tune cytokine mRNA stability and dampen inflammatory output. The overall profile, including terms like defense response to virus and cytokine-mediated signaling pathway, indicates that while immune functions are still engaged, they are recalibrated under FGA139 treatment. Rather than exacerbating inflammation like LPS, FGA139 appears to redirect microglial activity toward protective antiviral defense and immune regulation. These findings support the hypothesis that FGA139, a cysteine protease inhibitor, confers neuroprotective and anti-inflammatory effects, likely by attenuating excessive inflammatory cascades while maintaining immune competence through selective activation of interferon-related pathways. (**B**) Cellular components show enrichment in mitochondrial structures and membrane-bound organelles. “Mitochondrial envelope” shows the highest enrichment (strongest red color) with a log2 odds ratio around 5. Membrane-associated structures appear throughout the list, including “mitochondrial membrane” and “lysosomal membrane”. “Intracellular membrane-bounded organelle” has the largest dot size (indicating more genes) but appears less significant (purple). “Multivesicular body” shows a high log2 odds ratio but lower statistical significance (small blue dot). The strong enrichment in mitochondrial components indicates potential effects on energy metabolism and mitochondrial function. (**C**) For the Molecular Function category, the analysis revealed enrichment of terms associated with nucleic acid binding and immune signaling. Among the most prominent were STAT family protein binding (log odds ratio >7), adenylyltransferase activity, and double-stranded RNA binding (log odds ratios >5), pointing to enhanced transcriptional activation through the JAK-STAT pathway and increased antiviral surveillance mechanisms in FGA139-treated cells. *Y-axis: Lists Gene Ontology (GO) terms. X-axis (log2 Odds ratio): Represents the enrichment of each term. Higher values mean the term is more enriched (above than expected by chance). Dot color: reflects p-value significance according to the gradient. Dot size: reflects the number of genes involved (likely the count of genes associated with the term that are enriched)*.

## Supplementary figures

**Supplementary Table 1.**
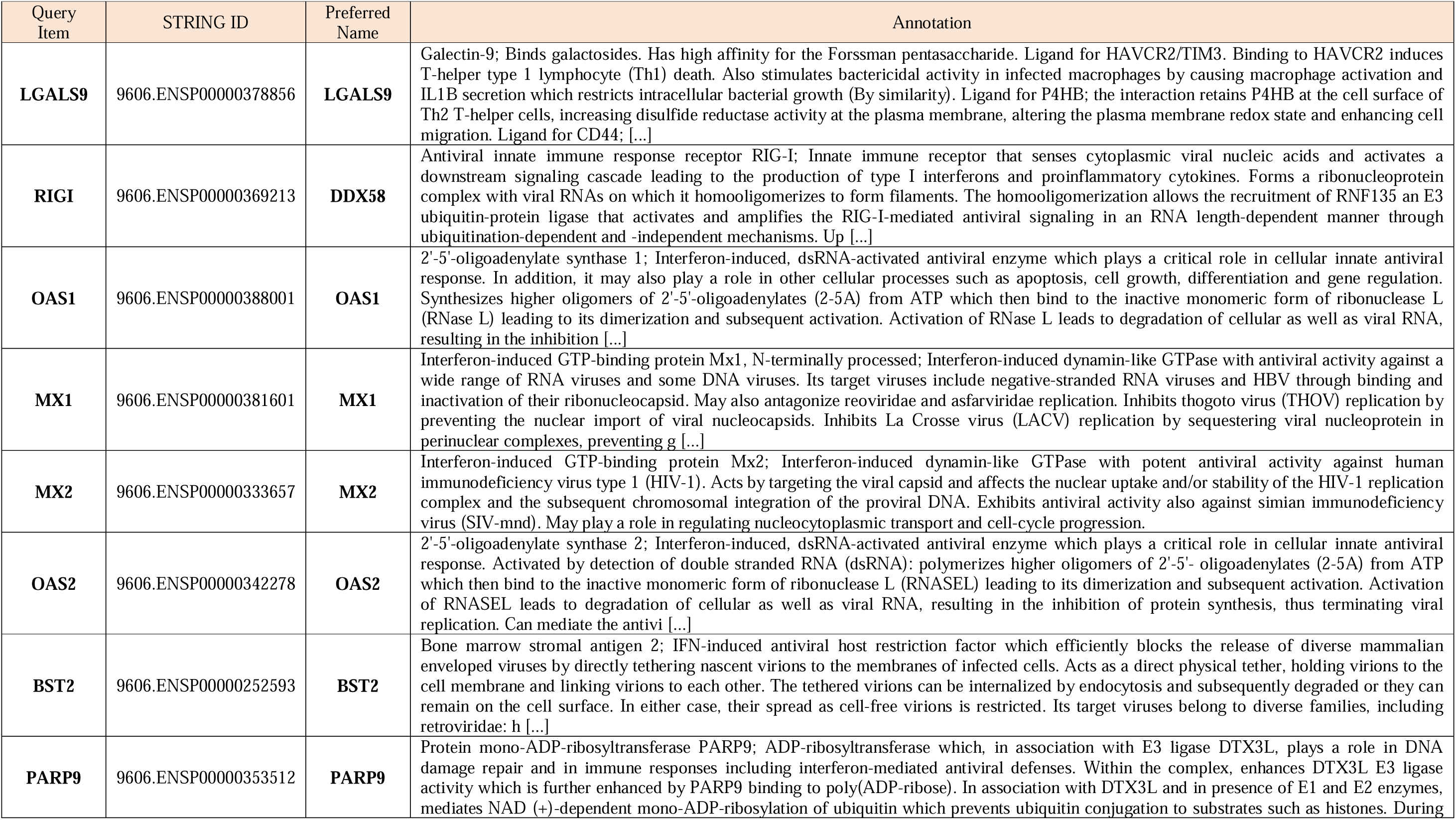
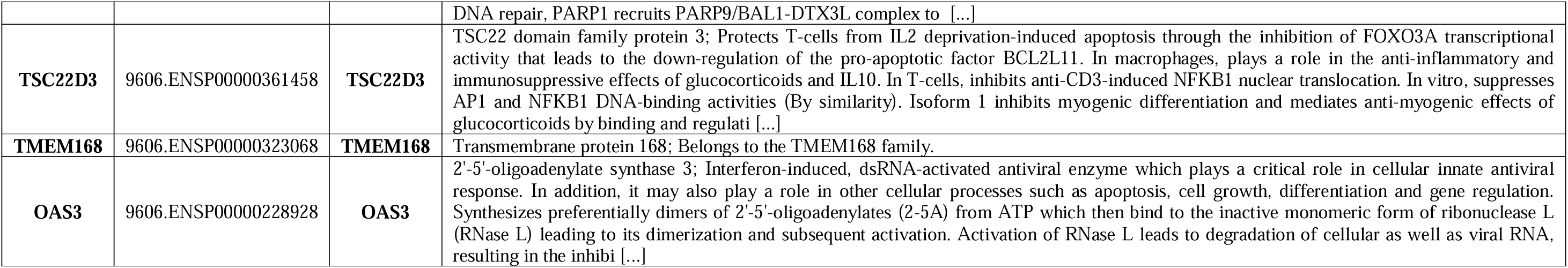
Functional characterization of differentially expressed genes in response to LPS versus NM treatment. Genes exhibiting significant expression changes (log fold change/ratios) were analyzed using STRING software. The table provides gene identifiers, STRING IDs, alternative names, and a summary of their main biological functions.

| Query Item | STRING ID | Preferred Name | Annotation |
| --- | --- | --- | --- |
| **LGALS9** | 9606.ENSP00000378856 | **LGALS9** | Galectin-9; Binds galactosides. Has high affinity for the Forssman pentasaccharide. Ligand for HAVCR2/TIM3. Binding to HAVCR2 induces T-helper type 1 lymphocyte (Th1) death. Also stimulates bactericidal activity in infected macrophages by causing macrophage activation and IL1B secretion which restricts intracellular bacterial growth (By similarity). Ligand for P4HB; the interaction retains P4HB at the cell surface of Th2 T-helper cells, increasing disulfide reductase activity at the plasma membrane, altering the plasma membrane redox state and enhancing cell migration. Ligand for CD44; [...] |
| **RIGI** | 9606.ENSP00000369213 | **DDX58** | Antiviral innate immune response receptor RIG-I; Innate immune receptor that senses cytoplasmic viral nucleic acids and activates a downstream signaling cascade leading to the production of type I interferons and proinflammatory cytokines. Forms a ribonucleoprotein complex with viral RNAs on which it homooligomerizes to form filaments. The homooligomerization allows the recruitment of RNF135 an E3 ubiquitin-protein ligase that activates and amplifies the RIG-I-mediated antiviral signaling in an RNA length-dependent manner through ubiquitination-dependent and -independent mechanisms. Up [...] |
| **OAS1** | 9606.ENSP00000388001 | **OAS1** | 2'-5'-oligoadenylate synthase 1; Interferon-induced, dsRNA-activated antiviral enzyme which plays a critical role in cellular innate antiviral response. In addition, it may also play a role in other cellular processes such as apoptosis, cell growth, differentiation and gene regulation. Synthesizes higher oligomers of 2'-5'-oligoadenylates (2-5A) from ATP which then bind to the inactive monomeric form of ribonuclease L (RNase L) leading to its dimerization and subsequent activation. Activation of RNase L leads to degradation of cellular as well as viral RNA, resulting in the inhibition [...] |
| **MX1** | 9606.ENSP00000381601 | **MX1** | Interferon-induced GTP-binding protein Mx1, N-terminally processed; Interferon-induced dynamin-like GTPase with antiviral activity against a wide range of RNA viruses and some DNA viruses. Its target viruses include negative-stranded RNA viruses and HBV through binding and inactivation of their ribonucleocapsid. May also antagonize reoviridae and asfarviridae replication. Inhibits togoto virus (THOV) replication by preventing the nuclear import of viral nucleocapsids. Inhibits La Crosse virus (LACV) replication by sequestering viral nucleoprotein in perinuclear complexes, preventing g [...] |
| **MX2** | 9606.ENSP00000333657 | **MX2** | Interferon-induced GTP-binding protein Mx2; Interferon-induced dynamin-like GTPase with potent antiviral activity against human immunodeficiency virus type 1 (HIV-1). Acts by targeting the viral capsid and affects the nuclear uptake and/or stability of the HIV-1 replication complex and the subsequent chromosomal integration of the proviral DNA. Exhibits antiviral activity also against simian immunodeficiency virus (SIV-mnd). May play a role in regulating nucleocytoplasmic transport and cell-cycle progression. |
| **OAS2** | 9606.ENSP00000342278 | **OAS2** | 2'-5'-oligoadenylate synthase 2; Interferon-induced, dsRNA-activated antiviral enzyme which plays a critical role in cellular innate antiviral response. Activated by detection of double stranded RNA (dsRNA); polymerizes higher oligomers of 2'-5'- oligoadenylates (2-5A) from ATP which then bind to the inactive monomeric form of ribonuclease L (RNASEL) leading to its dimerization and subsequent activation. Activation of RNASEL leads to degradation of cellular as well as viral RNA, resulting in the inhibition of protein synthesis, thus terminating viral replication. Can mediate the anti-viral response [...] |
| **BST2** | 9606.ENSP00000252593 | **BST2** | Bone marrow stromal antigen 2; IFN-induced antiviral host restriction factor which efficiently blocks the release of diverse mammalian enveloped viruses by directly tethering nascent virions to the membranes of infected cells. Acts as a direct physical tether, holding virions to the cell membrane and linking virions to each other. The tethered virions can be internalized by endocytosis and subsequently degraded or they can remain on the cell surface. In either case, their spread as cell-free virions is restricted. Its target viruses belong to diverse families, including retroviridae: h [...] |
| **PARP9** | 9606.ENSP00000353512 | **PARP9** | Protein mono-ADP-ribosyltransferase PARP9; ADP-ribosyltransferase which, in association with E3 ligase DTX3L, plays a role in DNA damage repair and in immune responses including interferon-mediated antiviral defenses. Within the complex, enhances DTX3L E3 ligase activity which is further enhanced by PARP9 binding to poly(ADP-ribose). In association with DTX3L and in presence of E1 and E2 enzymes, mediates NAD (+)-dependent mono-ADP-ribosylation of ubiquitin which prevents ubiquitin conjugation to substrates such as histones. During |
|  |  |  | DNA repair, PARP1 recruits PARP9/BAL1-DTX3L complex to [...] |
| **TSC22D3** | 9606.ENSP00000361458 | **TSC22D3** | TSC22 domain family protein 3; Protects T-cells from IL2 deprivation-induced apoptosis through the inhibition of FOXO3A transcriptional activity that leads to the down-regulation of the pro-apoptotic factor BCL2L11. In macrophages, plays a role in the anti-inflammatory and immunosuppressive effects of glucocorticoids and IL10. In T-cells, inhibits anti-CD3-induced NFkB1 nuclear translocation. In vitro, suppresses AP1 and NFkB1 DNA-binding activities (By similarity). Isoform 1 inhibits myogenic differentiation and mediates anti-myogenic effects of glucocorticoids by binding and regulati [...] |
| **TMEM168** | 9606.ENSP00000323068 | **TMEM168** | Transmembrane protein 168; Belongs to the TMEM168 family. |
| **OAS3** | 9606.ENSP00000228928 | **OAS3** | 2'-5'-oligoadenylate synthase 3; Interferon-induced, dsRNA-activated antiviral enzyme which plays a critical role in cellular innate antiviral response. In addition, it may also play a role in other cellular processes such as apoptosis, cell growth, differentiation and gene regulation. Synthesizes preferentially dimers of 2'-5'-oligoadenylates (2-5A) from ATP which then bind to the inactive monomeric form of ribonuclease L (RNase L) leading to its dimerization and subsequent activation. Activation of RNase L leads to degradation of cellular as well as viral RNA, resulting in the inhibi [...] |

**Supplementary table 2:**
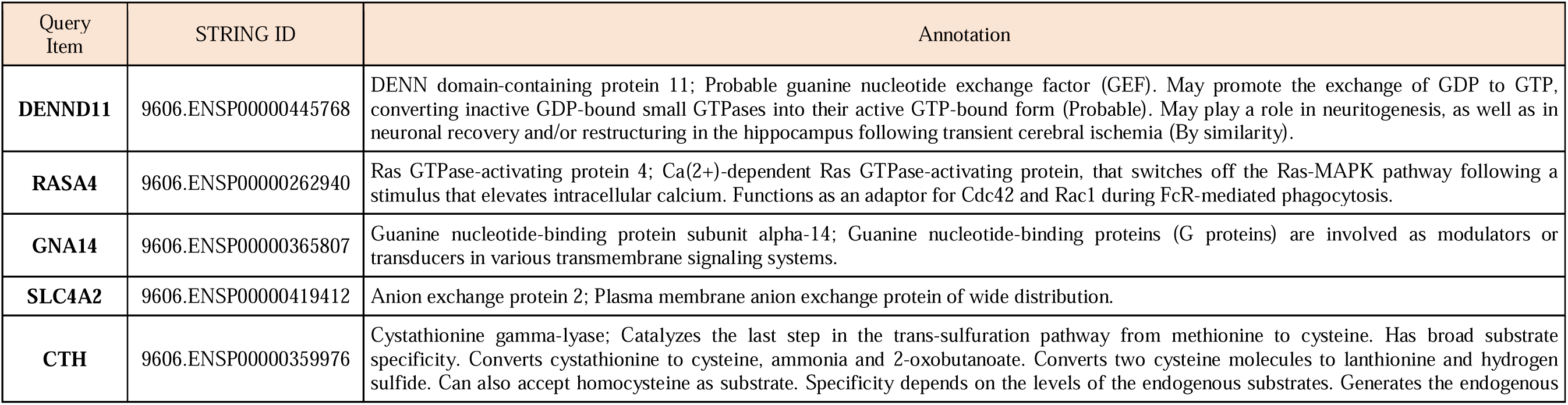
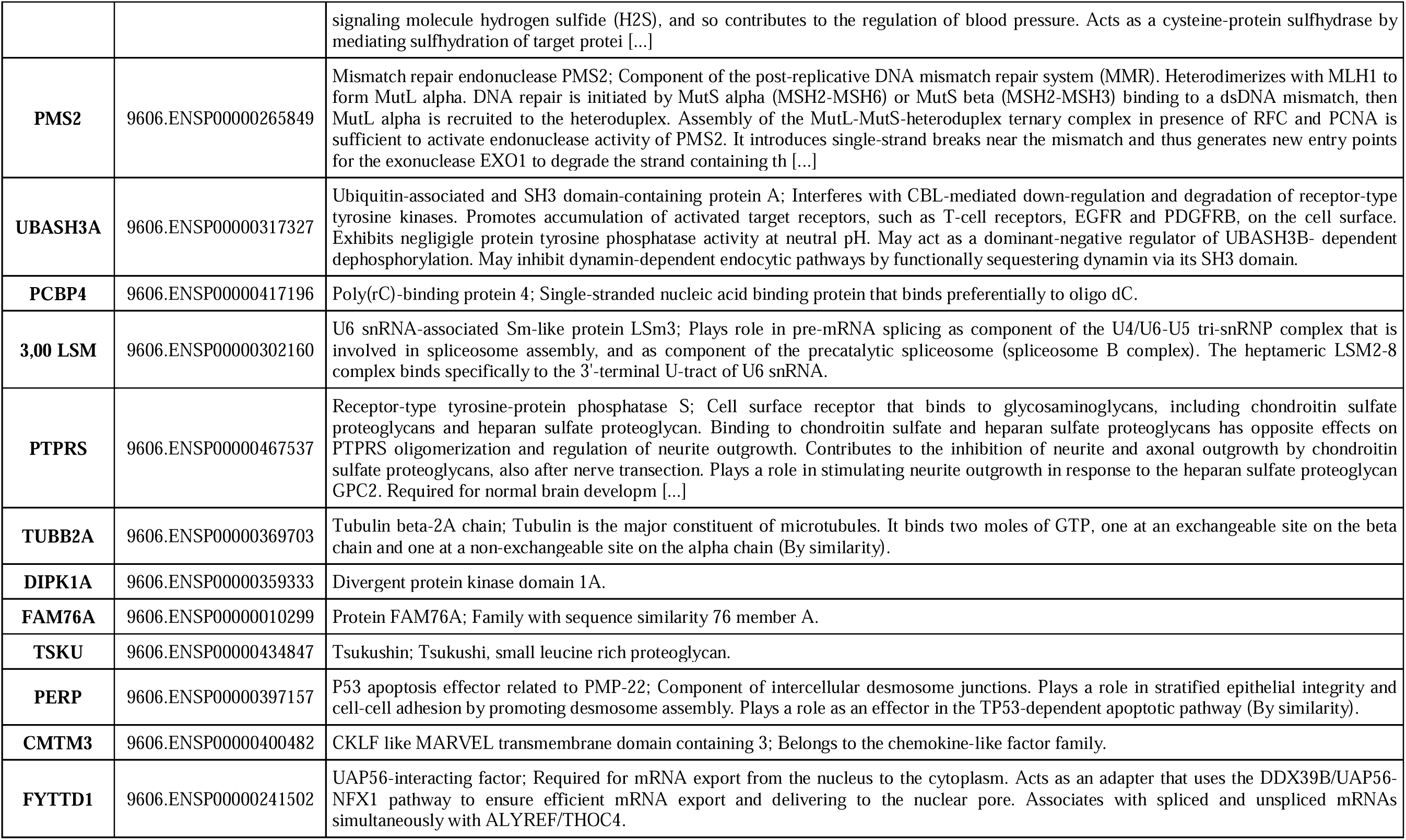
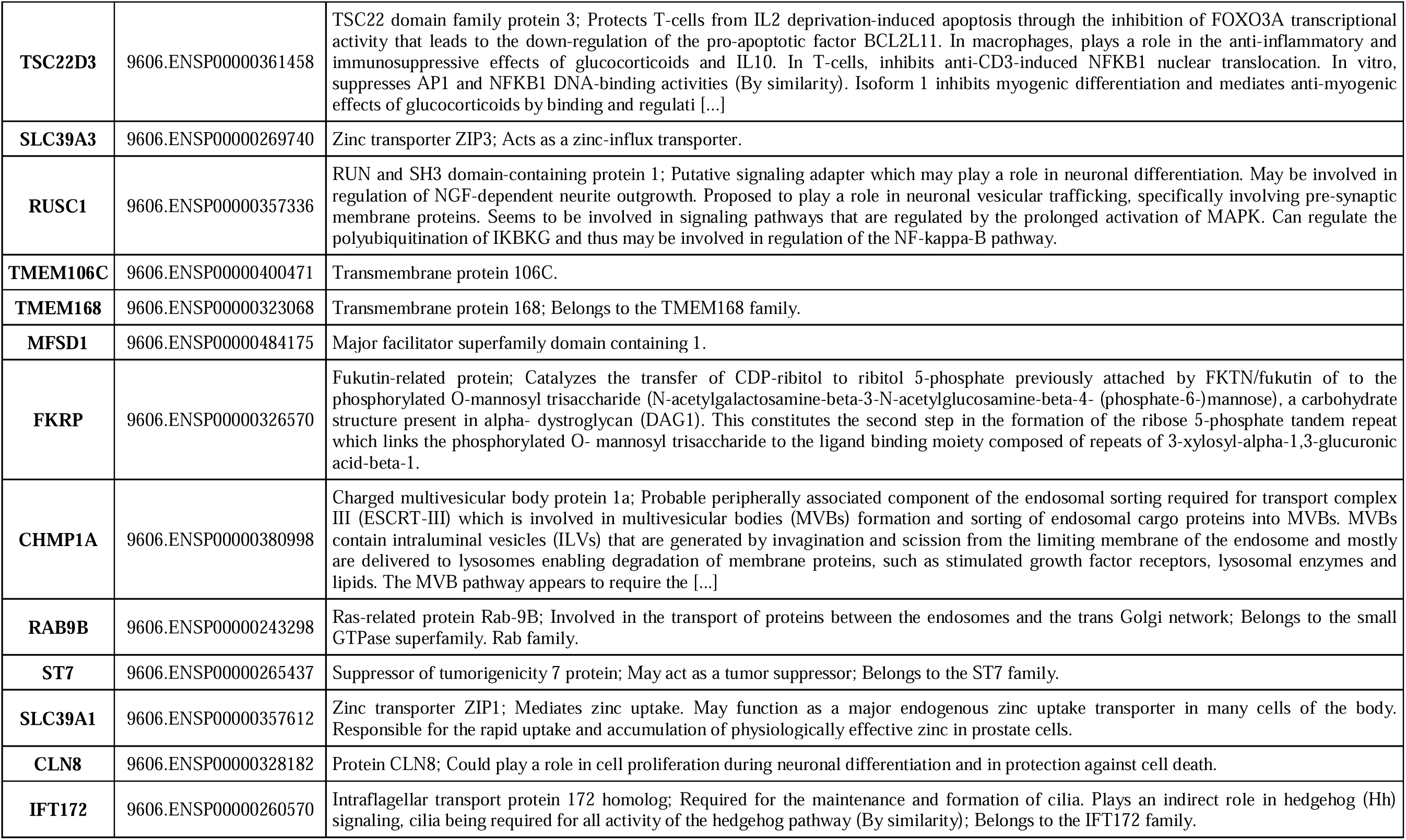
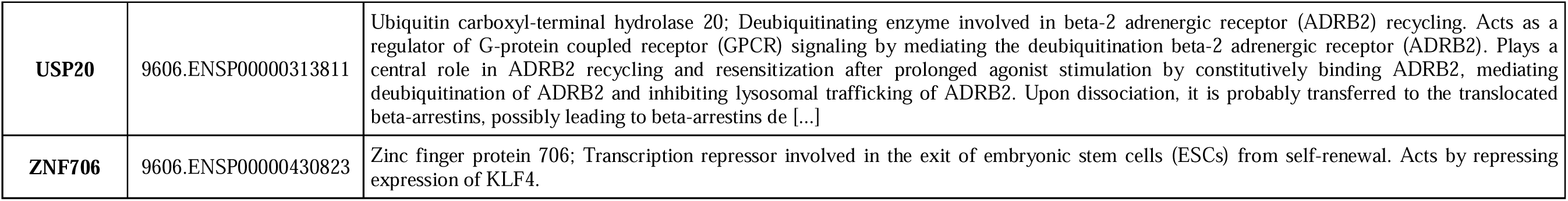
Functional characterization of differentially expressed genes in response to FGA139 + LPS vs LPS treatment. Genes exhibiting significant expression changes (log fold change/ratios) were analyzed using STRING software. The table provides gene identifiers, STRING IDs, alternative names, and a summary of their main biological functions.

